# A population code for spatial representation in the larval zebrafish telencephalon

**DOI:** 10.1101/2023.11.12.566708

**Authors:** Chuyu Yang, Lorenz Mammen, Byoungsoo Kim, Meng Li, Drew N Robson, Jennifer M Li

## Abstract

The vertebrate telencephalon is the site of complex cognitive processes, such as spatial cognition. The larval zebrafish telencephalon is a compact circuit of only ∼10,000 neurons that contains potentially homologous structures to the mammalian basal ganglia and limbic system (e.g., the hippocampus). However, despite long-standing evidence that spatial navigation and learning in zebrafish requires an intact telencephalon, cells believed to underlie spatial cognition in the mammalian hippocampus (e.g., place cells) have yet to be established in any fish species. Using a tracking microscope to image brain-wide activity at cellular resolution in freely swimming larval zebrafish, we compute the spatial information of neurons throughout the zebrafish brain. Strikingly, in every animal we recorded, cells with the highest spatial specificity are enriched in the zebrafish telencephalon. These cells form a population code of space, from which we can decode the animal’s spatial location across time. By continuous recording of population-level activity, we find that the activity manifold of place cells gradually untangles over time. Through systematic manipulation of allothetic and idiothetic cues, we demonstrate that place cells in the zebrafish telencephalon integrate multiple sources of information. By analysis of neighborhood distance between cells across environments, we find that the spatial representation in the zebrafish telencephalon partially generalizes across environments, suggesting that preconfigured network states may have been a feature of spatial computation that emerged early in vertebrate evolution.

## INTRODUCTION

Animals generate an internal map of their environment through spatial exploration^1–3^. A rich history of research in mammals has demonstrated that this spatial map is created from multiple sources of information, including distal and proximal landmarks, geometric features of the environment, and the animal’s integrated self-motion^4–8^. Over half a century of research in mammals has uncovered the key computational building blocks of spatial cognition, beginning with the seminal discovery of place cells in the mammalian hippocampus^1^, and expanding to grid cells in the entorhinal cortex^9^, border vector cells (BVCs) and heading direction cells in the subiculum^10–13^, and theta cells in the septum^14,15^. Much less is understood about when and how these building blocks emerged across evolution.

While behavioral studies suggest that spatial cognitive ability evolved early in evolution and is present in vertebrate species such as teleost fish^16^, previous studies have found no clear evidence for place cells outside of birds^17^ and mammals^1,18^. Instead, it has been proposed that teleost fish use an entirely distinct set of computational units (e.g., border cells but not place cells) for spatial cognition^19,20^. Indeed, the location or even existence of the teleost hippocampus is a matter of intense debate, with conflicting models that posit either that the hippocampal and paralimbic system encompasses most of the teleost telencephalon or that no hippocampal analog exists in the teleost brain^21,22^. This debate has been exceedingly difficult to resolve, due in large part to the lack of comprehensive functional data in the teleost brain during spatial navigation.

Previous studies have neither identified a comprehensive network of place cells that can be collectively used to decode an animal’s spatial location across time, nor produced definitive evidence that place cells do not exist in the teleost brain^19,20,23,24^. Several studies have found anecdotal evidence of neurons with spatial information in goldfish and electric fish. However, the goldfish recording identified just one cell with significant spatial information, in an animal that did not traverse the entire chamber^23^, and the spatially tuned neurons in electric fish are landmark cells, but not place cells^24^. More recent electrophysiological studies in goldfish have failed to discover any spatially tuned place cells^19,20^. However, given that the location of a hippocampal analog in the fish telencephalon is unclear, electrophysiological recording of a small number of neurons may limit the discovery of a comprehensive network of neurons that encode place.

In light of these open questions, the larval zebrafish represent an ideal model system, where brain wide imaging is possible at cellular resolution^25,26^. To date, however, most neural imaging studies in larval zebrafish have used tethered preparations that prevent naturalistic spatial navigation. It is unclear to what extent tethered larval zebrafish can truly undergo self-initiated spatial exploration, rather than reflexively responding to visual cues (e.g., optomotor response^26^, phototaxis^27^, prey detection^28,29^). Furthermore, freely swimming larval zebrafish spontaneously generate movement bouts at around 1 Hz during exploration, while tethered larval zebrafish generate tail flicks at only < 0.1 Hz^30^. Mammalian studies have shown that place cell activity is most robust during continuous running^31^. Thus, even if a tethered animal can be shown to navigate in virtual reality, its suppressed movement rate makes it potentially difficult for the discovery of place cells.

To overcome these limitations, we performed brain-wide imaging in freely swimming larval zebrafish^30,32^ to comprehensively characterize the functional properties and anatomical distribution of spatial information in the teleost brain. In contrast to cells that encode heading and speed, which are both enriched in the fish rhombencephalon^30,33,34^, cells that encode spatial position are primarily concentrated in the telencephalon. Collectively, these place-encoding neurons can be used to decode the animal’s spatial location across time. By projecting this population code onto a 2D activity manifold, we observe untangling of spatial representation across time, suggesting experience-dependent refinement of place codes in the zebrafish brain.

Through an extensive series of environmental manipulations, we find that this network of place-encoding neurons potentially integrates both allothetic and idiothetic information, akin to mammalian place cells. Finally, these place-encoding neurons show evidence of assembly structure, in support of recent work in mammals demonstrating stronger correlations between place cell pairs than previously expected^35–37^, which together suggest that preconfigured place cell networks may have been a feature of spatial computation that emerged early in vertebrate evolution. The existence of place-encoding neurons in the larval zebrafish telencephalon represents a significant step forward in our current understanding of the possible building blocks of spatial cognition in the fish brain, the functional role of early vertebrate telencephalon, and the evolutionary origin of spatial cognition.

## RESULTS

### Identification of cells encoding positional information in the larval zebrafish brain

To determine whether cells in the larval zebrafish brain encode spatial information, we constructed a rectangular behavioral chamber with landmark cues (**Fig. 1a**, **Methods**). The chamber was 50 mm × 25 mm and contained landmark cues at two opposing geometric corners (cue A and cue B), whereas the two remaining corners were unmarked. A transparent 1.5 mm wide inner wall ran along the entire perimeter of the chamber, preventing the animal from directly contacting the landmarks. The entire behavioral chamber was illuminated with diffuse white light. Larval zebrafish expressing pan-neuronal H2B-GCaMP6s (6-8 days post fertilization, 4-5 mm in length) were placed into this chamber and permitted to freely explore the entire chamber for 90 min (**Fig. 1b**).

**Figure 1.**
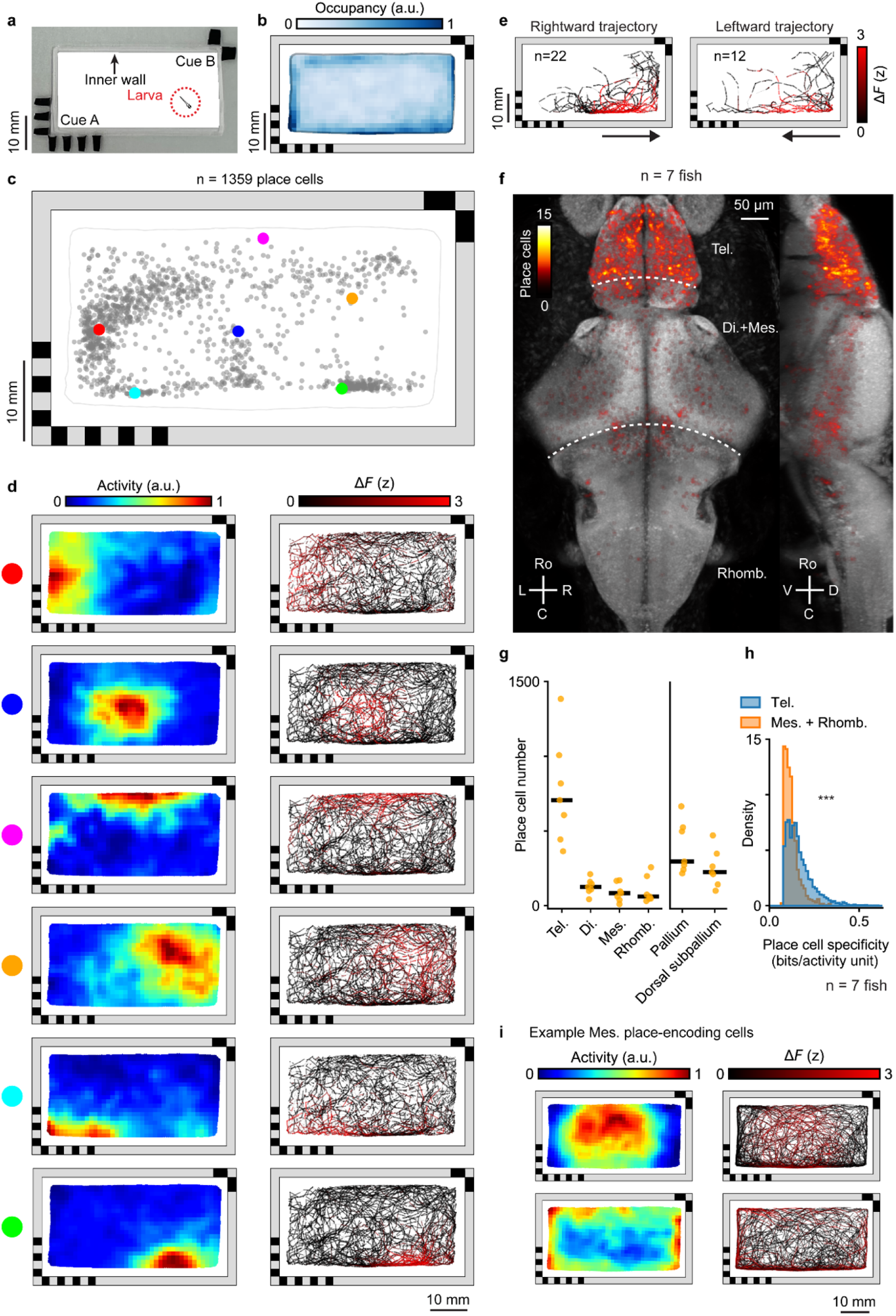
Behavioral arena with visual cues reveals place cells in the larval zebrafish brain. **a**, Rectangular behavioral arena with two landmark cues containing different visual patterns on opposing corners. There is a transparent inner PMDS wall (1.5 mm wide) preventing the animal from directly contacting the landmarks. **b**, Normalized median chamber occupancy across 7 animals. **c**, Distribution of place field center of mass for all place cells with a confined place field in an example fish (**Methods**). Each dot represents the center of mass of one cell’s place field (**Methods**). The six example cells (colored dots) are shown in **d**. **d**, Spatial activity maps (left, averaged neural response by location, **Methods**) and corresponding animal trajectories (right, color-coded by neural activity) are shown for six example cells from **c**. **e**, Responses of an example cell with a non-directional place field for two orientations of traversals (leftwards and rightwards) through the place field. Traversals across the 90 min experiment are color-coded by the neural activity. Orientation is classified by the animal’s heading as it enters the place field. **f**, Anatomical distribution of place cells across 7 animals plotted on the reference brain (maximum projection, **Methods**). **g**, Number of place cells in each brain region. Each animal is shown individually (yellow dots, n = 7 animals) along with the median across animals (black line). Anatomical subdivisions of the telencephalon (pallium and dorsal subpallium, **Extended Data Fig. 2d**) are plotted separately. Tel., telencephalon; Mes., mesencephalon; Di., diencephalon; Rhomb., rhombencephalon. **h**, Comparison of spatial specificity for place cells found in the telencephalon and place cells found in the mesencephalon and rhombencephalon (n = 7 animals, p < 10^−5^, one-sided Mann– Whitney U test). **i,** Spatial activity maps and animal trajectories for two example mesencephalic cells with significant spatial information encoding the interior (top) or periphery of the chamber (bottom).

During these 90 min of spatial exploration, we recorded each animal’s brain-wide calcium activity at cellular resolution with an imaging rate of 2 volumes/s, and then extracted the fluorescence across time *F*(*t*) for each neuronal centroid in the brain using non-negative matrix factorization (NMF)^38^ (**Methods**). Based on the baseline-corrected neural activity Δ*F*(*t*) and the position of the animal in space, we generated a 2D spatial activity map for each cell, calculated its spatial information content normalized by its mean activity (i.e., spatial specificity)^39^, and identified the center of mass (COM) location of its place field (**Fig. 1c-d**) (see **Methods** for detailed description of each analysis).

To identify cells with significant spatial specificity, we shuffled the activity of each cell 1000 times by circular permutation. We generated an associated 2D spatial activity map and computed the spatial specificity of each shuffled activity trace. The shuffled data forms a null distribution of spatial specificity for a given cell. A cell was designated as place-encoding (or “place cell”) if its spatial specificity was significantly greater than 1) the null distribution generated from its shuffled activity and 2) the distribution of spatial specificity across all cells (**Methods**). Using our significance criteria, we identified 1081 ± 329 (mean ± s.d.) place-encoding cells or “place cells” in each animal, with place fields distributed throughout the chamber (**Fig. 1c-d, Extended Data Fig. 1a**).

Using linear regression, we independently confirm that the activity of the identified place cells is primarily predicted by the animal’s position in space, and not by other behavioral parameters such as heading or speed (**Extended Data Fig. 1b**, **Methods**). Spatial position also accounts for a greater portion of activity variance in place cells compared with other cells in the telencephalon (p < 10^−5^, one-sided Mann–Whitney U test, **Extended Data Fig. 1c**). Thus, despite large differences in visual experience as the animal traverses the same location in space with distinct heading direction, we can identify cells that consistently encode a preferred spatial location (**Fig. 1e**). The stability of spatial representation for a given cell can be further quantified by measuring the spatial correlation between two halves of the experiment. Place cells have an average spatial correlation of 0.61 ± 0.18 (median ± s.d.) across the 90 min experiment and are consistently activated on multiple traversals through the same place field (**Extended Data Fig. 1d-e, Supplementary Video 1**). We note that slow GCaMP decay kinetics combined with the speed of animal motion likely blur the spatial activity map of a given place cell, resulting in place fields that appear larger than their true size (**Extended Data Fig. 1f-h, Methods**). Based on the behavioral data (**Extended Data Fig. 1f**) and calcium indicator kinetics, which depend on the firing rate^40^, we estimate the radius of the place fields to be spatially blurred by 0.17 to 1.31 mm (**Methods**).

### Place cells are enriched in the telencephalon of larval zebrafish

The anatomical distribution of spatial information in the brain is remarkably consistent across animals. We find that in every animal, the telencephalon contains the highest number and fraction of place cells (**Fig. 1f-g, Extended Data Fig. 2a-f**) regardless of the chamber geometry or landmarks (**Extended Data Fig. 2c**). This enrichment of place cells in the telencephalon can also be observed by simply ranking cells across the brain according to their spatial specificity (**Extended Data Fig. 2g**) or spatial information (**Extended Data Fig. 2h**). In every animal, the cells with highest ranked spatial specificity are located primarily in the telencephalon (91 ± 9 %, mean ± s.d., for the top 100 cells).

Place cells that encode a given spatial location are not clustered together anatomically (**Extended Data Fig. 2j**). Across place cell pairs, we find little correlation between their anatomical distance and the similarity of their spatial activity maps (**Extended Data Fig. 2k**).

Outside the telencephalon, cells with the highest spatial specificity are located in a small number of bilaterally symmetric neuronal clusters near the boundary between the mesencephalon and rhombencephalon (**Fig. 1f**). However, the spatial specificity or spatial information of these cells in the mesencephalon or rhombencephalon is significantly lower (p < 10^−5^, one-sided Mann–Whitney U test) than those within the telencephalon (**Fig. 1h, Extended Data Fig. 2i**). The place fields show larger activation areas, representing either the interior or the border of the chamber (**Fig. 1i**). These non-telencephalic place cells appear to contain coarse-grained spatial information, while the telencephalon contains finer-scale spatial information that forms a population code of space.

### Majority of spatially specific cells in the telencephalon are not consistent with the BVC model

A previous study has proposed that teleost fish may use border vector cells (BVCs) instead of place cells as the primary computational units for spatial cognition^19^. BVCs are neurons that fire at a given distance and allocentric heading from a boundary^10,11,13^ (**Extended Data Fig. 3a**), rather than at a localized position in space. Thus, the spatial activity of BVCs contain two unique properties – 1) BVCs fire along an entire boundary at a given allocentric heading and 2) when borders are duplicated, BVCs also duplicate their place fields. To test for the presence or absence of border vector cells, a movable wall was inserted and removed in the middle of an extended rectangular chamber across three sessions (**Extended Data Fig. 3b, Methods**). For all candidate telencephalic neurons with a preferred firing orientation towards the left or right side of the chamber (1922 ± 298 cells, mean ± s.d.), the BVC model predicts duplication of firing fields in response to wall insertion. We fit the activity of each candidate neuron with the BVC model (**Extended Data Fig. 3c, Methods**), and find that only a small percentage (1.60 ± 1.56%, mean ± s.d.) of neurons in the telencephalon fit the BVC model (prediction performance > 0.6, Pearson correlation between the predicted change in activity and the observed change in activity) (**Extended Data Fig. 3d, Methods**). The majority of place cells do not show duplication of place fields and thus do not fit the BVC model (**Extended Data Fig. 3e**). The mean prediction performance of the BVC model across all place cells at the left and right edges of the chamber is low, with a Pearson correlation of 0.05 ± 0.3 (mean ± s.d.).

### Telencephalic place cells form a population code of space

Collectively, place cells in the telencephalon span spatial locations throughout the entire behavioral chamber, with high variance in the degree of spatial representation of each location across animals (**Fig. 2a**, **Extended Data Fig. 4a-b**). The relative density of spatial representation does not correlate with the animal’s behavioral occupancy in the chamber (**Extended Data Fig. 4c-d**).

**Figure 2.**
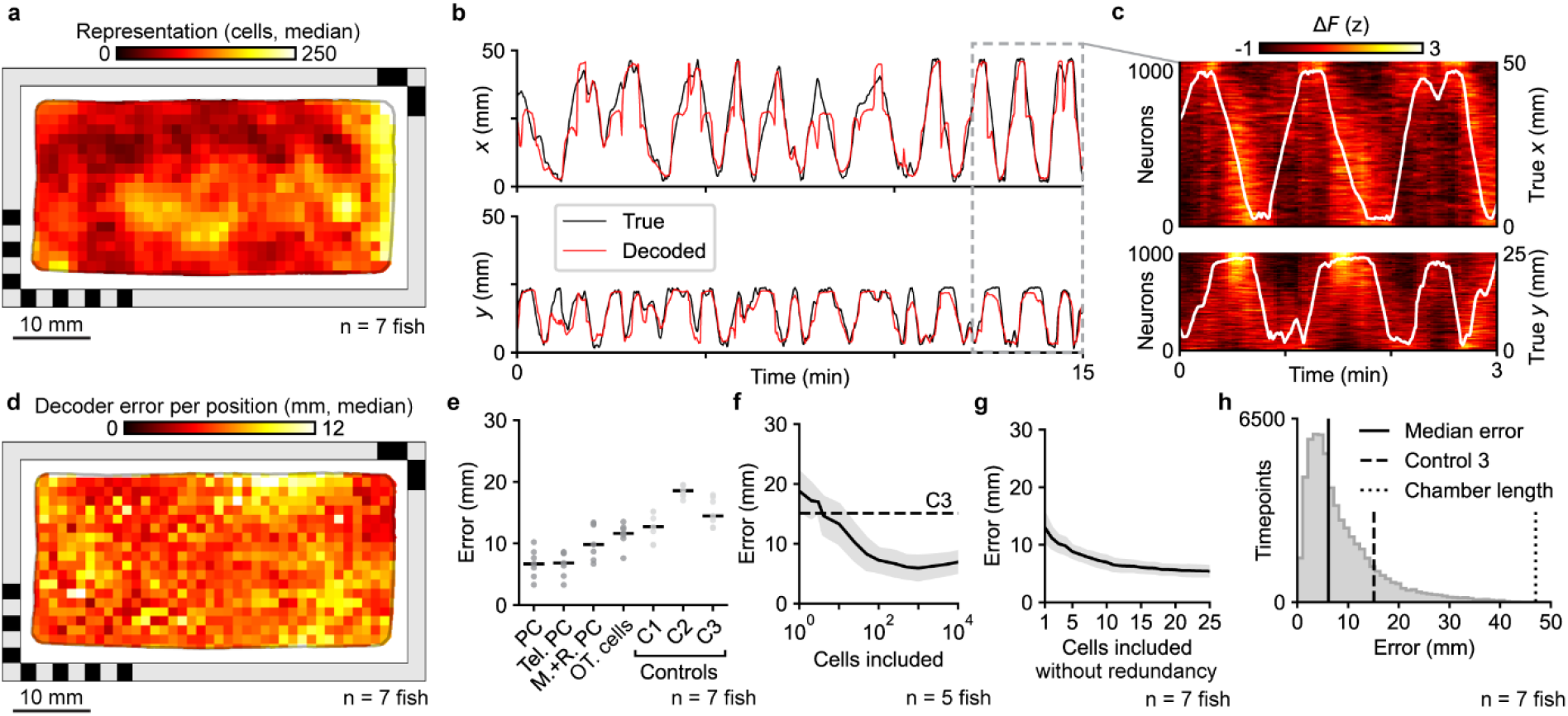
Physical location of the larval zebrafish can be decoded from the population activity of place cells. **a,** Distribution of telencephalic place fields encoding different locations in the behavioral arena (n = 7 animals, median across animals is shown). To quantify the density of the place fields across space, each spatial activity map was binarized (**Methods**). **b,** To decode the physical location of the animal, a direct basis decoder with no free parameters was applied to the 1000 telencephalic place cells with highest spatial specificity (**Methods**). Example traces show the *x* (top) and *y* (bottom) coordinates of the true (black) and decoded (red) animal location. **c**, Activity of place cells across the last three minutes of **b**, together with true *x* (top) and *y* (bottom) coordinates of the animal (white overlay). Activity is shown two ways, either sorting each cell by the *x* component (top) or *y* component (bottom) of its place field’s center of mass. **d,** Median decoder error at each binned animal location within the behavioral arena (n = 7 animals). **e,** Decoding performance from 1000 cells with the highest spatial tuning within different brain regions: place cells (PC) across the entire brain, telencephalic place cells (Tel. PC), mes- and rhombencephalic place cells (M.+R. PC), and cells from the optic tectum (OT. cells). Three controls are shown: average error of a decoder based on random non-place cells (C1), from a decoder that outputs chamber positions uniformly at random (C2), and from a decoder that invariantly outputs the centroid of the animal’s spatial occupancy map (C3). Black horizontal bars represent the median across 7 animals. **f,** Decoder error as a function of the number of telencephalic cells included, in descending order of spatial tuning (black, mean; gray, std. dev.). Cells were first ranked by spatial specificity, and the number of input cells to the decoder was increased by gradually relaxing the threshold for spatial tuning (**Methods**, n = 5 animals with ≥ 10 000 telencephalic cells, 2 animals with < 10,000 recorded cells in the telencephalon were not included in this analysis). **g,** Decoder error as a function of the number of place cells included, using greedy selection to minimize redundancy. Cells were iteratively selected based on which cell improves the decoding result most (black, mean; gray, std. dev.). **h,** Distribution of decoder error across time (pooled, n = 7 animals). Vertical lines indicate the median error (6.11 mm, black line), behavior-informed baseline (15.08 mm, dashed line), and accessible length of the chamber long axis (47 mm, dotted line). For all panels except **f** and **g**, the 1000 cells with the highest spatial specificity were used as input to the decoder (n = 7 animals).

To determine whether this population code of neural activity can be used to decode the animal’s physical position within the chamber, we adapted the direct basis decoder^41^ for spatial location. This decoder has no free parameters and predicts the animal’s position at a given time by summing all the spatial activity maps, each weighted by the corresponding cell’s activity at that time. To avoid circularity, decoding was only performed on held out timepoints not included in spatial activity map construction (**Fig. 2, Extended Data Fig. 5, Methods**).

This linear decoder is able to predict the animal’s spatial location to within 6.69 ± 2.34 mm (median ± s.d.) using the cells with the highest spatial specificity (up to the top 1000) throughout the whole brain. (**Fig. 2b-e, h**, **Extended Data Fig. 5a**). Using only the activity of spatially tuned cells cells in the telencephalon, the decoder predicts spatial position with a similar degree of accuracy, at an error of 6.82 ± 2.05 mm (median ± s.d.), which significantly outperforms decoding from spatially tuned cells in the mesencephalon and rhombencephalon (error of 9.84 ± 2.89 mm, median ± s.d.), neurons in the optic tectum (11.63 ± 1.89 mm, median ± s.d.), random cells across the brain (error of 12.70 ± 2.30 mm, median ± s.d.), and the baseline decoding error of 15.08 ± 2.30 mm (median ± s.d., **Fig. 2e**, **Methods**). The optimal time lag for the decoder is 2.5 s (**Extended Data Fig. 5c**). To determine how many place cells are needed to achieve optimal decoding, we ranked cells by their degree of spatial tuning and varied the number of input cells (**Methods**). The decoder reached peak performance (5.93 ± 1.98, median ± s.d.) with the 1000 cells with highest spatial tuning (**Fig. 2f**).

However, since the top 1000 spatially selective neurons may contain redundant encodings of position, we additionally demonstrate that, by selecting place cells with non-redundant place fields, a decoding accuracy of 5.41 ± 0.81 mm (mean ± s.d.) can be achieved with a minimal set of 25 place cells (**Fig. 2g**, **Methods**). In fact, the non-redundant decoder starts to outperform the decoder using all cells from 17 cells onwards. The worse performance of the decoder using all cells is a consequence of the simple linear decoder design, which values the information of all cells equally. Hence, less specific neurons can influence the decoder and slightly worsen the results. The advantages of this decoder, however, are the limited assumptions of the model, easy interpretability, as well as no possibility of overfitting.

### Telencephalic place cells form a spatial activity manifold that untangles over time

We find subtle changes in the spatial representation of a stable environment across a 90 min period (**Fig. 3**). We define an early and late period as the first and last 30 min of the session, respectively. To control for potential differences in spatial coverage between the two periods, we subsampled time points in both early and late periods to completely equalize their spatial coverage (**Methods**). When the population-level activity is embedded in a 2D isomap^42^, the spatial manifold becomes more untangled across time (**Fig. 3a**, **Supplementary Video 2**). To quantify this, we define a metric, neighbor distance (**Fig. 3b**), that identifies for each timepoint *t_i_* the 30 nearest timepoints in the embedded 2D manifold space and measures their average distance to *t_i_* in physical space. From the early to late period, the distance relationships in the activity manifold increasingly resemble the distance relationships in the physical chamber, resulting in a significant decrease in neighborhood distance (p < 10^−4^ for 6/7 animals, each tested by one-sided Mann-Whitney U test, **Fig. 3b**). Neighborhood distance computed directly from neural activity without dimensionality reduction also shows a similar decrease in neighborhood distance across time (p < 10^−5^ for 6/7 animals, one-sided Mann-Whitney U test).

**Figure 3.**
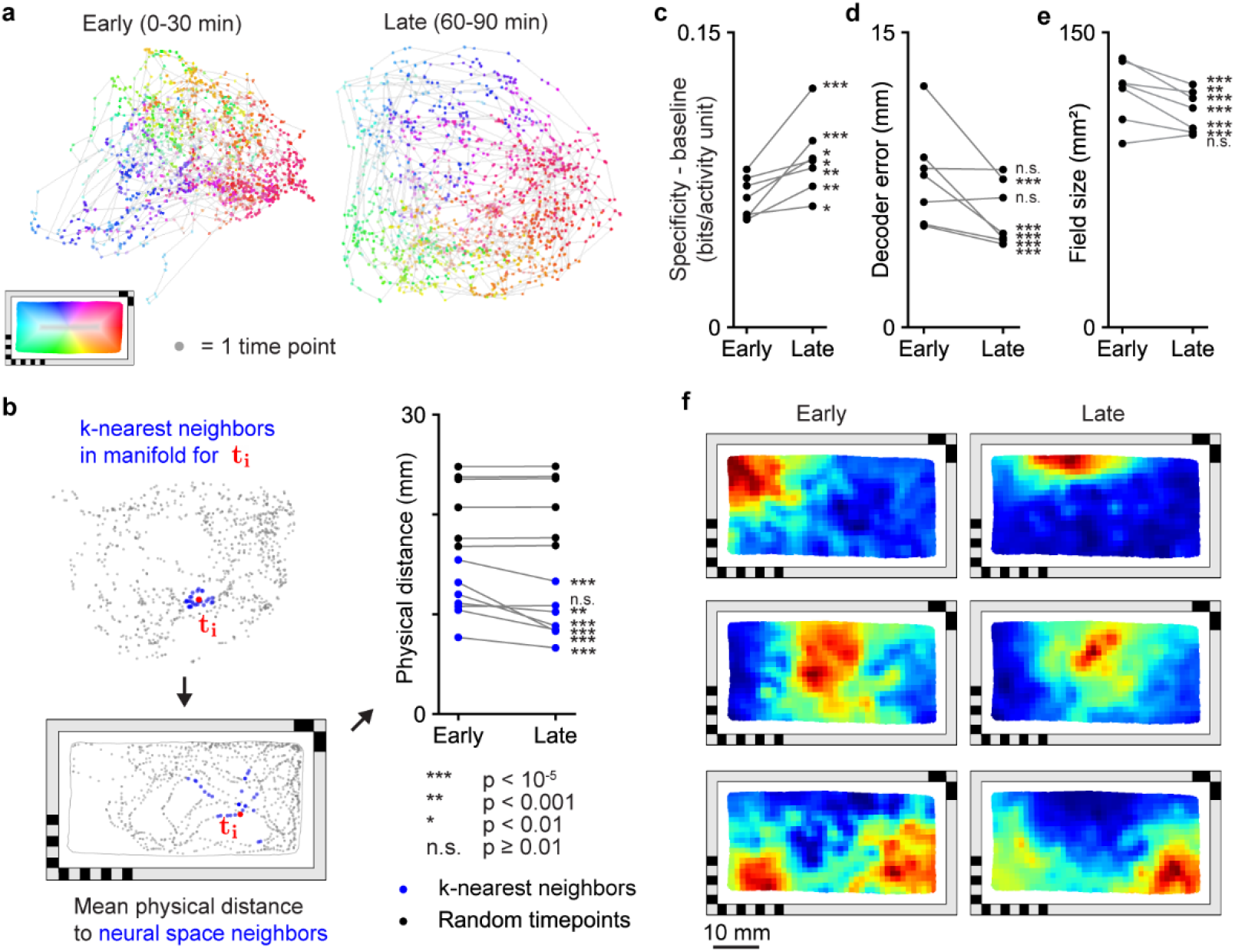
Activity manifold of place cells untangles spatial representation across time. **a**, Change in manifold structure across two stages of the experiment (early, 0-30 min; late, 60-90 min). Individual time points are color-coded by the animal’s location within the behavioral arena. For the whole figure, telencephalic place cells that are used are defined across the experiment, excluding the first and last 15 minutes, and behavioral coverage was equalized between the two examined time intervals before comparisons are made (**Methods**). **b**, The mean physical distance to 30 neighboring points in the 2D manifold space, averaged across all time points (left). Change of this distance between the early (0-30 min) and late (60-90 min) stages of the experiment (blue) and a baseline (black, 30 randomly selected timepoints instead of neighbors, **Methods**). Each animal is shown individually, and changes in the neighbor distance within animals are tested using a one-sided Mann-Whitney U test. **c**, Change of mean spatial specificity across telencephalic place cells from the first 30 minutes (early stage) to the last 30 minutes (late stage) of the experiment. Mean spatial specificity of place cells is baseline-corrected by subtracting the mean spatial specificity of 100 random cells. Each animal is shown individually and the baseline-subtracted specificity distributions between early and late periods are compared using a one-sided Wilcoxon signed-rank test. **d**, Change of mean decoder error from the first 30 minutes (early) to the last 30 minutes (late) of the experiment. Each animal is shown individually and changes in decoder error within animals are tested using a one-sided Mann-Whitney U test. **e**, Change of place field size from the first 30 minutes (early) to the last 30 minutes (late) of the experiment. Each animal is shown individually, and changes in place field size within animals are tested using a one-sided Wilcoxon signed-rank test. **f**, Example spatial activity maps for early and late stages of the experiment. Each row is one place cell. In the whole figure, significance is abbreviated by *** for p < 10^−5^, ** for p < 0.001, * for p < 0.01, n.s. for p ≥ 0.01.

The untangling of the spatial activity manifold is accompanied by a significant increase in spatial specificity (p < 0.01 for 7/7 animals, per-animal one-sided Wilcoxon signed-rank test, **Fig. 3c**), decrease in decoding error (p < 10^−5^ for 5/7 animals, per-animal one-sided Mann-Whitney U test, **Fig. 3d**), and decrease in place field size (p < 10^−4^ for 6/7 animals, per-animal Wilcoxon signed-rank test, **Fig. 3e**). The untangling of the activity manifold, along with the increased spatial specificity of individual cells (see **Fig. 3f** for example spatial activity maps), together suggest refinement of the population code for spatial representation across time.

### Integration of idiothetic and allocentric information in spatial representation

A hallmark of mammalian place cells is their ability to integrate information from both idiothetic and allocentric cues. Idiothetic information enables the animal to update its position in space based on path integration, while an allocentric reference frame (i.e. formed from a combination of landmarks and geometric boundaries) orients, constrains, and stabilizes the animal’s internal position estimate. In the face of changing environmental conditions, this multimodal computation leads to complex changes in spatial tuning^8,43^, with idiothetic information typically dominating in initially novel environments and allothetic information becoming the primary source of information in overly familiarized environments^8,44^.

In the following sections, we created a new set of behavioral paradigms for larval zebrafish, in which we systematically manipulated allothetic information while either maintaining or interrupting idiothetic information. We find evidence that both idiothetic and allothetic information are essential to the spatial tuning properties of place cells in the telencephalon (**Fig. 4-6, Extended Data Fig. 6-7**). First, without disrupting path integration (i.e. the animal is not removed from the chamber between sessions), spatial tuning is largely maintained despite changes in landmarks, environmental boundary and visual illumination (**Fig. 4, Extended Data Fig. 7**). Second, when path integration is interrupted (i.e. when the animal is physically removed from the chamber between sessions), the animal’s previous spatial activity map can be recovered based on partial or complete allocentric information (e.g., the presence of consistent landmarks and/or geometric features) (**Fig. 5a-l**). When both geometric and landmark cues are significantly altered, however, a new spatial activity map is generated (**Fig. 5m-p**). We will elaborate on each case below.

**Figure 4.**
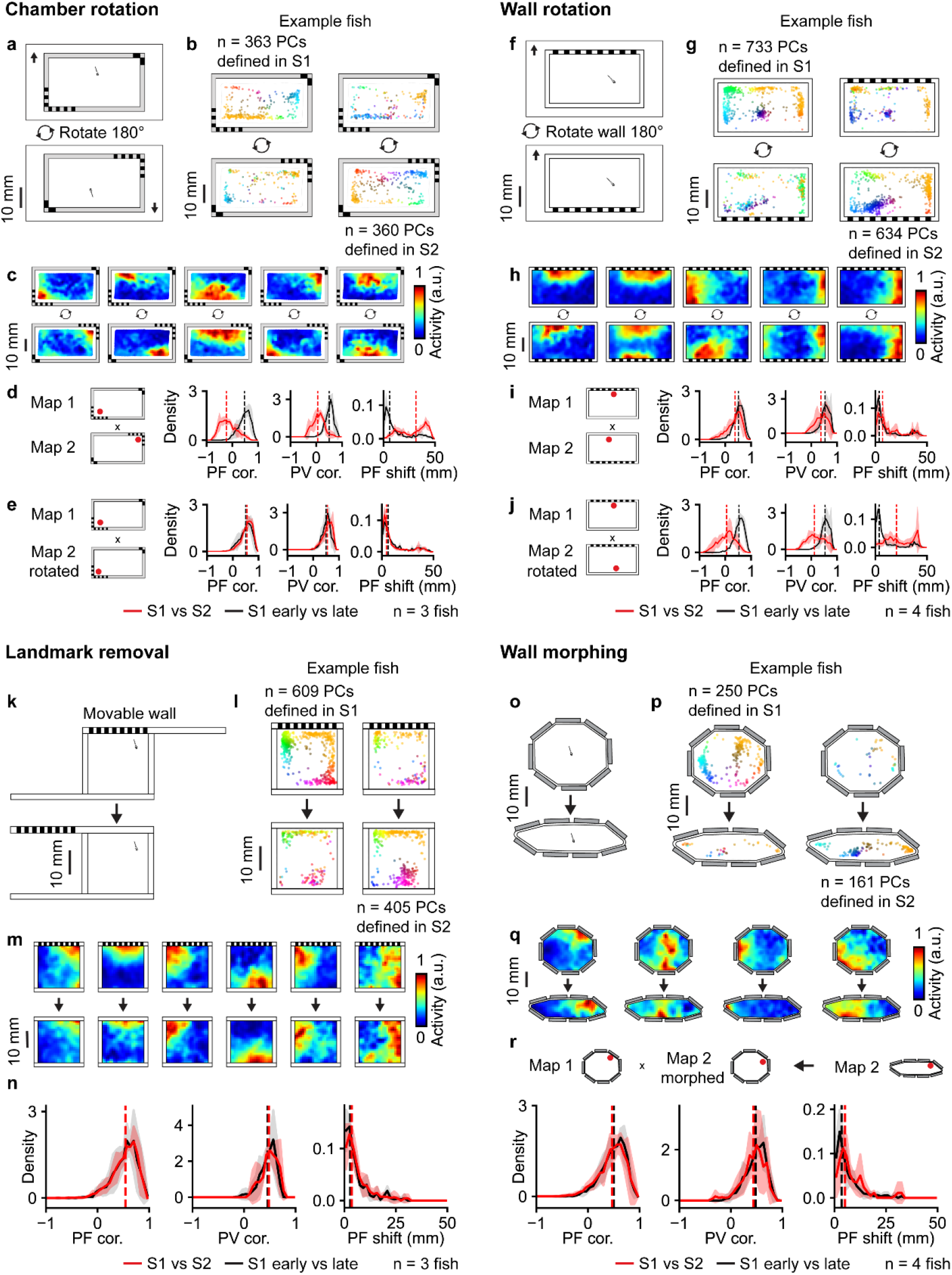
Environmental manipulation provides evidence for the involvement of path integration in place cell activity. **a**, Schematic of chamber rotation experiment. Between the 1^st^ and the 2^nd^ recording sessions, the entire chamber including the fish is rotated 180 degrees relative to the microscope (**Methods**). **b**, Change in the distribution of telencephalic place cells with confined place fields, plotted separately for telencephalic place cells defined in the 1^st^ session (left) or defined in the 2^nd^ session (right) for an example animal. The top row shows the distribution of place fields in the 1^st^ session, and bottom row shows the distribution of place fields for the 2^nd^ session. Each dot represents the center of mass of one cell’s place field, colored by its position in the chamber. **c**, Example spatial activity maps before and after the chamber rotation. Each column is one cell. **d-e**, Summary of the changes in telencephalic place cells across the two sessions by comparing the maps either directly in the microscope reference frame (**d**) or in the chamber reference frame (**e**) by rotating the spatial activity maps (**Methods**). From left to right, schematic of the map comparison, place field correlation (PF cor.), population vector correlation (PV cor.), and shift in the place field location (PF shift, **Methods**). All telencephalic cells identified as place cells in either session were used. Comparisons between the 1^st^ and 2^nd^ sessions (red) are plotted together with a reference distribution, obtained by comparing the early (1^st^ half) and late (2^nd^ half) stages of the 1^st^ session (black). Solid lines are the mean distributions across all fish, and the shaded region indicates the standard deviation across all fish. Vertical dashed lines are the medians of the averaged distributions. **f**, Schematic of wall rotation experiment. In comparison to the experiment in **a**, here only the chamber wall, but not the fish, is rotated 180 degrees relative to both the fish and the microscope (**Methods**). **g-j**, Analysis of wall rotation experiment as in **b-e**. **k**, Schematic of landmark removal experiment. A grating pattern is painted on a movable wall, which was remotely controlled by magnets to become invisible to the fish in the 2^nd^ recording session (**Methods**). **l-n**, Analysis of landmark removal experiment as in **b-d**. **o**, Schematic of wall morphing experiment. Between the 1^st^ and the 2^nd^ recording sessions, the chamber comprised of 8 hinged pieces of stainless steel was morphed into a different shape (**Methods**). **p-q**, Analysis of wall morphing experiment as in **b-c. r**, similar to **e**, but the registration is done in a nonrigid way by mapping pixels from the 2^nd^ session to the 1^st^ session according to their distance to the wall anchor points (**Methods**). By morphing the 2^nd^ session spatial activity map, we represented the activity of both sessions in terms of the spatial bins of session 1, facilitating comparisons. The results of statistical tests (one-sided Wilcoxon signed-rank test comparing experimental and control distributions for each animal) for PF correlation, PV correlation, and PF shift in each experimental condition are summarized in **Fig. 6**.

**Figure 5.**
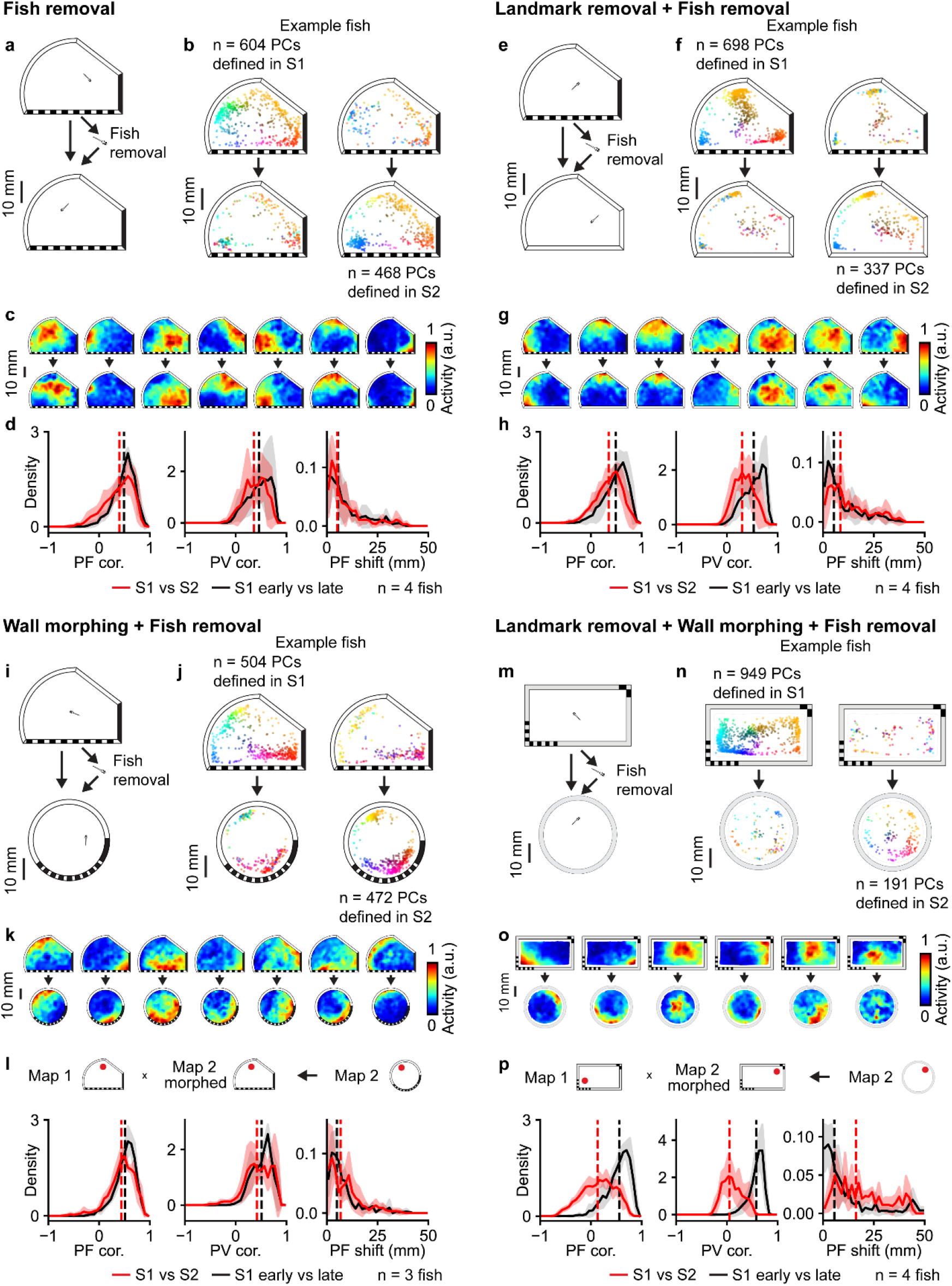
Manipulation of environmental geometry and landmarks after breaking path integration reveals their relationship to place cell activity. **a**, Schematic of the fish removal experiment. The fish was removed from the chamber to break path integration between recording sessions performed in the same asymmetric chamber with landmarks (**Methods**). **b-d**, Analysis of fish removal experiments as in **Fig. 4b-d**. **e**, Schematic of the landmark removal experiment with fish removal. After the 1^st^ recording session in an asymmetric chamber with landmarks, the fish was removed and transferred into a chamber with the same geometry but no landmarks for the 2^nd^ recording session (**Methods**). **f-h**, Analysis of landmark removal experiment with fish removal as in **Fig. 4b-d**. **i.** Schematic of the wall morphing experiment with fish removal. After the 1^st^ recording session in an asymmetric chamber with landmarks, the fish was removed and transferred into a chamber with the same landmarks but a different geometry for the 2^nd^ recording session (**Methods**). **j-l**, Analysis of wall morphing experiment with fish removal as in **Fig. 4p-r**. **m**, Schematic of the experiment combining landmark removal, wall morphing, and fish removal. After the 1^st^ recording session, the fish was removed and transferred into a chamber with a different geometry and no landmarks for the 2^nd^ recording session (**Methods**). **n-p**, Analysis of experiment combining landmark removal, wall morphing, and fish removal as in **Fig. 4p-r**. The results of statistical tests (one-sided Wilcoxon signed-rank test comparing experimental and control distributions for each animal) for PF correlation, PV correlation, and PF shift in each experimental condition are summarized in **Fig. 6**.

**Figure 6.**
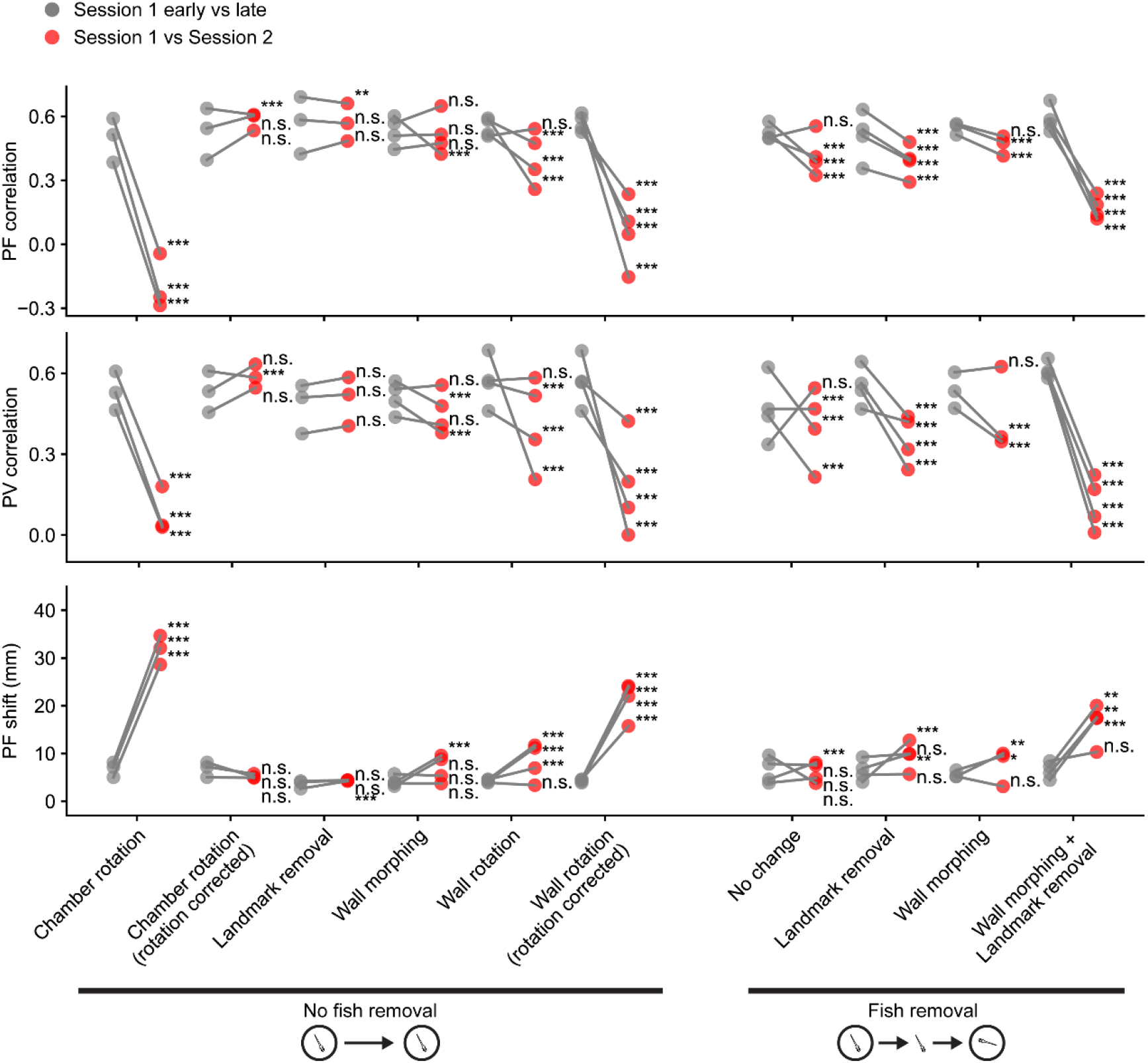
Summary of spatial activity map changes of telencephalic place cells under different environmental manipulations. The median of PF correlation (top), PV correlation (mid), and PF shift (bottom) for individual fish. Gray dots are comparisons between the early (1^st^ half) and late (2^nd^ half) stages of the 1^st^ session. The red dots are comparisons made between the 1^st^ session and the 2^nd^ session. Experiments from **Fig. 4** and **Fig. 5** are represented on the horizontal axis: chamber rotation (**Fig. 4a**), landmark removal (**Fig. 4k**), wall morphing (**Fig. 4o**), wall rotation (**Fig. 4f**), fish removal with no change in the chamber (**Fig. 5a**), landmark removal with fish removal (**Fig. 5e**), wall morphing with fish removal (**Fig. 5i**), and wall morphing with landmark removal and fish removal (**Fig. 5m**). Data from the same fish are connected by gray lines. The results of statistical tests are marked for each fish separately (one-sided Wilcoxon signed-rank test, *** for p < 10^−5^, ** for p < 0.001, * for p < 0.01, n.s. for p ≥ 0.01).

### Minimal remapping is observed when path integration is uninterrupted

In principle, three sources of information could contribute to the spatial activity maps observed in the telencephalon – allothetic information external to the behavioral chamber (e.g., from the imaging room or microscope), allothetic information internal to the behavioral chamber (e.g., distal landmarks on the chamber walls and chamber geometry), and idiothetic information generated by the movement of the animal itself.

First, to test whether allothetic information external to the behavioral chamber could affect the spatial turning properties of the place cells, we performed a simple rotation experiment. After an initial imaging session of 90 min (session 1), we rotated the chamber 180°, and then resumed recording of neural activity throughout the brain for an additional 60 min (session 2) (**Fig. 4a, Methods**). This manipulation only alters the animal’s spatial relationship to allothetic information external to the behavioral chamber, while allothetic information specific to the behavioral chamber (i.e. distal landmarks on the chamber walls and chamber geometry), as well as idiothetic information (i.e. path integration) are preserved.

Post-rotation, we find that the spatial activity map rotated 180°, corresponding to the chamber rotation (**Fig. 4b-c**). To quantify the degree of change in spatial representation, we compute three metrics – 1) place field (PF) correlation, which is the correlation of spatial activity maps between sessions, 2) PF shift, which is the change in the center of mass (COM) of place fields across sessions, and 3) population vector (PV) correlation, which is the average correlation between population activity vectors at each spatial location (**Methods**). Direct comparison of the spatial activity map across the two sessions without adjusting for the chamber rotation results in a PF correlation of −0.23 (with an interquartile range, IQR, of −0.44 to 0.03, defined as median of quartiles 1 and 3, **Methods**), PV correlation of 0.06 (IQR −0.1 to 0.2), and PF shift of 31.47 mm (IQR 15.59 to 39.49 mm), all of which are significantly different from a control comparison of early and late periods within session 1 (**Fig. 4d**, **Fig. 6**, p < 10^−5^ for 3/3 fish, per-fish one-sided Wilcoxon signed-rank test).

In contrast, after applying a 180° rotation to the spatial activity map in session 2, PF correlation increased to 0.55 (IQR 0.39 to 0.68), PV correlation increased to 0.55 (IQR 0.41 to 0.66), and PF shift decreased to 4.52 mm (IQR 1.87 to 12.10 mm), all of which became indistinguishable from internal control comparisons (**Fig. 4e**, **Fig. 6**, n.s., 3/3 fish for PF shift; n.s., 2/3 fish for PF and PV correlation, n.s. defined as p ≥ 0.01, per-fish one-sided Wilcoxon signed-rank test). Overall, these results suggest that allothetic information external to the chamber does not contribute significantly to the spatial tuning properties of these cells, while allothetic information specific to the behavioral chamber and idiothetic information are sufficient to stabilize the spatial activity map across sessions.

Next, we asked whether allocentric information specific to the chamber or idiothetic information contributes more to the spatial tuning properties of place cells. To test this, we created a chamber wall that can be remotely rotated without removing the fish or disassembling the water-sealed chamber (**Methods** on chamber construction and experiment design). First, we recorded neural and behavioral activity for 45-90 min (session 1) within a rectangular chamber with a unique landmark on one wall (**Fig. 4f**). We then remotely rotated the chamber wall 180° without removing the fish. The animal was recorded for an additional 45-75 min (session 2).

If idiothetic information dominates over allothetic information, then the spatial activity map in session 2 should directly correspond to the spatial activity map in session 1. If the converse is true, then the spatial activity map in session 2 should rotate 180°. We find that the resulting spatial activity map in session 2 is intermediate between these two extremes (**Fig. 4g-j**). A control comparison of early and late periods within session 1 results in a PF correlation of 0.53 (IQR 0.39 to 0.65), PV correlation of 0.53 (IQR 0.42 to 0.64), and PF shift of 3.55 mm (IQR 1.55 to 8.37 mm). In contrast to controls, direct comparison of spatial activity maps between session 1 and 2 show a significantly lower PF correlation of 0.38 (IQR 0.15 to 0.53) and PV correlation of 0.37 (IQR 0.18 to 0.53), and a significantly higher PF shift of 6.85 mm (IQR 3.12 to 17.48). The comparison of spatial activity maps after 180° rotation of the spatial activity map in session 2 results in an even lower PF correlation of 0.05 (IQR −0.2 to 0.25) and PV correlation of 0.12 (IQR −0.1 to 0.39), and even higher PF shift of 20.47 mm (IQR 12.31 to 36.23). Thus, compared with the control, we find significant remapping based on PF correlation, PV correlation, and PF shift using both direct (p < 10^−5^, 3/4 fish, per-fish one-sided Wilcoxon signed-rank test) and rotated session 2 spatial activity maps (p < 10^−5^, 4/4 fish, per-fish one-sided Wilcoxon signed-rank test, **Fig. 4i-j**, **Fig. 6**). However, the spatial activity map in session 2 shows greater direct (rather than 180° rotated) correspondence to the spatial activity map in session 1, suggesting that idiothetic information is dominant over allothetic information when the two information sources come into conflict.

To further establish that idiothetic information alone can contribute significantly to spatial tuning, we performed two classic experiments – 1) transition from light to dark^45^, and 2) removing landmarks from the environment^8,46^. In the first case, we recorded neural activity and behavior for 60 min under bright diffuse white light illumination (7.05 µW/mm^2^) in a circular chamber (session 1), followed by 60 min with no white light illumination in session 2 (**Extended Data Fig. 7a, Methods**). We note that a localized blue excitation spot (0.5 mm in radius) centered on the fish brain cannot be removed in these experiments due to its necessity for neural imaging (**Methods**). The animal was not removed from the chamber across the two recording sessions (i.e. path integration was not interrupted). We find that place fields are retained despite significant changes in visual illumination (**Extended Data Fig. 7b-c**), as shown by a PF correlation of 0.66 (IQR 0.51 to 0.77), a PV correlation of 0.58 (IQR 0.45 to 0.68), and a PF shift of 3.38 mm (IQR 1.50 to 6.79 mm), which are not significantly different from internal control comparisons (**Extended Data Fig. 7d-e**, n.s., 3/3 fish for PF correlation and PF shift; n.s., 2/3 fish for PV correlation, per-fish one-sided Wilcoxon signed-rank test).

In the second experiment, we created a movable wall that can be remotely positioned (**Fig. 4k**, 70 × 2 × 0.5 mm). Visible landmarks were added to the middle of the movable wall in session 1 (75-90 min). The wall was then moved to hide the landmarks in session 2 (60-100 min). The animal was not removed from the chamber between the two recording sessions (i.e. path integration was not interrupted). We find that place fields are retained despite landmark removal (**Fig. 4l-m**), as shown by a PF correlation of 0.54 (IQR 0.36 to 0.67), a PV correlation of 0.48 (IQR 0.37 to 0.58), and a PF shift of 3.61 mm (IQR 1.45 to 7.48 mm), which are comparable to controls with a PF correlation of 0.54 (IQR 0.36 to 0.67), a PV correlation of 0.46 (IQR 0.35 to 0.55), and a PF shift of 2.90 mm (IQR 1.01 to 6.46 mm) (**Fig. 4n**, **Fig. 6**, n.s., 3/3 fish for PV correlation; n.s., 2/3 fish for PF correlation and PF shift, per-fish one-sided Wilcoxon signed-rank test). We did not observe greater PF shift for cells with place fields near landmarks (**Extended Data Fig. 8**).

Lastly, we asked whether the retention of spatial tuning properties can be explained by the interaction of idiothetic cues and boundary conditions, as previously observed in mammals^47^. To test this, we created a morph chamber with eight flexibly connected segments that can be remotely adjusted without removing the animal (**Methods, Fig. 4o**). Indeed, we find that place fields mostly maintain their relative positions to each other across sessions (**Fig. 4p-r**, but not their absolute position in the chamber (**Extended Data Fig. 6a**). Direct comparison of PF shift across the two sessions is 8.41 mm (IQR 4.29 to 12.98 mm), which is significantly higher than the control PF shift of 3.44 mm (IQR 1.49 to 7.38 mm, p < 10^−5^, 3/4 fish, p < 0.01, 1/4 fish, per-fish one-sided Wilcoxon signed-rank test), confirming changes in the absolute positions of the place fields. Assuming no systematic angular change across sessions, we then performed a nonrigid transformation of the session 2 spatial activity maps to align the maps across sessions (**Methods**). PF shift across sessions decreased to 5.13 mm (IQR 2.75 to 10.11 mm), (**Fig. 4r**, **Fig. 6**). PF correlation and PV correlation are 0.47 (IQR 0.29 to 0.62) and 0.44 (IQR 0.29 to 0.56), respectively, which are similar to the control PF correlation of 0.5 (IQR 0.32 to 0.63) and control PV correlation of 0.48 (IQR 0.33 to 0.6) (**Fig. 4r**, **Fig. 6**).

### Recovery of spatial activity maps based on allocentric information

Next, we ask the converse question of whether, after interruption of path integration (e.g., by removing the animal from the chamber), a larval zebrafish can recover its spatial activity map based on allothetic cues (e.g., when returned to the same chamber). To do this, we recorded an animal for 60-90 min in a geometrically asymmetric behavioral chamber with distinct landmarks (session 1). The animal was then physically removed from the chamber (**Methods**), set aside for several minutes, returned to the chamber at a different location and heading from where it was removed, and then imaged for another 45-100 min (session 2, **Fig. 5a**). To control for any potential olfactory cues, the chamber was either thoroughly cleaned with soap and isopropanol between sessions (n = 2 fish, **Extended Data Fig. 6f**) or the bottom of the chamber was rotated after the animal was removed from the chamber (n = 2 fish, **Extended Data Fig. 6g**). In both cases, we find that the spatial activity map was maintained between session 1 and 2 despite the interruption in path integration and scrambling of any potential olfactory cues (**Fig. 5b-c**), as shown by a PF correlation of 0.40 (IQR 0.17 to 0.57), a PV correlation of 0.36 (IQR 0.19 to 0.51), and a PF shift of 5.05 mm (IQR 2.13 to 11.76 mm). PF shift was not significantly different from controls for most animals (n.s., 3/4 fish, per-fish one-sided Wilcoxon signed-rank test, **Fig. 5d**), suggesting that place field locations are largely maintained across the two sessions. PF and PV correlation were moderately lower than controls (**Fig. 5d**, **Fig. 6**, p < 10^−5^, 3/4 fish, n.s., 1/4 fish for PF correlation; p < 10^−5^, 3/4 fish, n.s., 1/4 fish for PV correlation; per-fish one-sided Wilcoxon signed-rank test), suggesting that the precise shape and activity level of place fields partially changed.

Furthermore, similar to mammals^36,43^, we find that even partial allocentric information is sufficient for significant recovery of the spatial activity map. We demonstrate this with partial removal of landmarks without changing environmental geometry (**Fig. 5e-g**) or changing environmental geometry while maintaining the same landmarks (**Fig. 5i-k**). In both cases, we recorded the animal for 60-90 min in session 1, removed the animal from the chamber and placed the animal into a new chamber with altered landmarks or geometry, and then recorded the animal in the new chamber for session 2 (45-90 min). Chambers are thoroughly cleaned between sessions (**Methods**).

For landmark removal, we find a PF correlation of 0.35 (IQR 0.16 to 0.50), a PV correlation of 0.30 (IQR 0.16 to 0.44), and a PF shift of 8.58 mm (IQR 4.17 to 17.79 mm, **Fig. 5h**), which suggests significant partial recovery of the spatial activity map compared to controls, which had a PF correlation of 0.49 (IQR 0.31 to 0.63), a PV correlation of 0.53 (IQR 0.34 to 0.66), and a PF shift of 5.38 mm (IQR 2.29 to 13.02 mm, **Fig. 5h**, **Fig. 6**, p < 10^−5^, 4/4 fish for PF correlation and PV correlation; p < 10^−5^, 1/4 fish, p < 0.001, 1/4 fish, n.s., 2/4 fish for PF shift; per-fish one-sided Wilcoxon signed-rank test).

In the case of geometric change, we performed a simple nonrigid transformation of the spatial activity maps in session 2 to align the maps across sessions (**Fig. 5l, Methods**), resulting in a PF correlation of 0.44 (IQR 0.27 to 0.59), a PV correlation of 0.42 (IQR 0.24 to 0.62), and a PF shift of 6.77 mm (IQR 2.21 to 11.69 mm). In comparison to controls, which had a PF correlation of 0.51 (IQR 0.38 to 0.62), a PV correlation of 0.51 (IQR 0.34 to 0.62), and a PF shift of 4.98 mm (IQR 2.35 to 9.04 mm), this suggests significant partial recovery of the spatial activity map (**Fig. 5l**, **Fig. 6**, p < 10^−5^, 2/3 fish, n.s.,1/3 fish for PF correlation; p < 10^−5^, 2/3 fish, n.s.,1/3 fish for PV correlation; p < 0.001, 1/3 fish, p < 0.01, 1/3 fish, n.s., 1/3 fish for PF shift, per-fish one-sided Wilcoxon signed-rank test).

Finally, when both chamber geometry and landmarks are altered (**Fig. 5m**), we find significantly greater remapping (**Fig. 5n-p**). We recorded the animal in session 1 (90 min), removed the animal from the chamber and placed the animal into a new chamber with distinct geometry and no landmarks on the walls, and then recorded session 2 (60-80 min). A simple nonrigid transformation was performed on the spatial activity maps in session 2 to align the maps across sessions (**Fig. 5p, Methods**). After applying the nonrigid transformation, we find a PF correlation of 0.13 (IQR −0.13 to 0.36), a PV correlation of 0.05 (IQR −0.07 to 0.21), and a PF shift of 16.13 mm (IQR 7.96 to 26.83 mm), all of which are significantly different from controls, which had a PF correlation of 0.56 (IQR 0.39 to 0.68), a PV correlation of 0.58 (IQR 0.48 to 0.65), and a PF shift of 5.49 mm (IQR 2.03 to 12.46 mm, **Fig. 5p**, **Fig. 6**, p < 10^−5^, 4/4 fish for PF correlation and PV correlation; p < 10^−5^, 1/4 fish, p < 0.001, 2/4 fish, n.s., 1/4 fish for PF shift; per-fish one-sided Wilcoxon signed-rank test).

The PF correlation, PV correlation and PF shift for each fish (shown as the median of each distribution) across all experiments are summarized in **Fig. 6**. The full distributions of PF correlation, PV correlation, and PF shift are summarized in **Extended Data Fig. 6** for three forms of cross-session comparisons of spatial activity maps – 1) direct comparison without registration (i.e. without applying a transformation to align maps across sessions, **Extended Data Fig. 6a**), 2) comparison after aligning landmarks across sessions and applying a nonrigid transformation when needed to correct for changes chamber geometry (**Extended Data Fig. 6b**, **Methods**), and 3) comparison of spatial activity maps after applying the best rotation angle that optimizes PF correlation between sessions and a nonrigid transformation when needed to correct for changes in chamber geometry (**Extended Data Fig. 6c-e**, **Methods**). The PF correlation for different possible rotation angles is shown in **Extended Data Fig. 6d**.

### Evidence for latent ensemble structure across environments

Studies in mammals have suggested animals may not employ entirely independent spatial maps despite significant changes in environmental landmarks and geometry, especially when recordings take place in the same room^36,48^. Different subregions of the hippocampus (e.g. CA1 and CA3) also show distinct remapping characteristics^48,49^. Even in the case of significant remapping, pairwise relationships between place cells can be partially maintained^35,37,43^. Finally, a recent study has shown that animals can use similar maps that generalize across environments with distinct geometry and landmarks^36^. However, this generalization can be difficult to detect due to coherent place field rotations that lead to misalignment of the spatial maps^36^.

When both environmental geometry and landmarks are altered, and path integration is interrupted, we find PF and PV correlation near 0 across sessions (**Fig. 5p**). Using these experiments, we asked whether there are latent local or global spatial relationships that are maintained across sessions despite significant remapping. First, we assessed whether local neighborhood relationships between place cells are retained by computing the percentage of neurons that remain neighbors across sessions (**Extended Data Fig. 9**, **Methods**). To avoid the possibility of confounds due to overlapping fluorescent signal between adjacent cells, we restricted the analysis to pairs of cells with a minimum anatomical distance of > 20 µm. In all animals, the neighbor retention percentages are significantly lower than controls (early vs. late period of the same session, p < 10^−5^, 4/4 fish, one-sided Wilcoxon signed-rank test), suggesting a significant disruption of neighborhood relationships during remapping. However, the percentage of neurons that remain neighbors are also significantly higher than random chance (**Extended Data Fig. 9**, p < 10^−5^, 4/4 fish, mean percentage of remaining neighbors compared with 1000 random shuffles, **Methods**), suggesting that local relationships between neurons are partially maintained.

Secondly, we ask whether there are latent global relationships between the spatial activity maps (e.g., coherent place field rotations^36^). We synthetically applied 72 possible rotations (in 5° increments, **Methods**) to the spatial activity map in session 2, and then used a nonrigid transformation to align the spatial activity maps across sessions. We identified the rotation that maximized the correlation between the two spatial activity maps, and found this to be variable across the 4 animals (131°, −147°, −86°, −100°). Application of the optimal rotation and nonrigid transformation of the session 2 spatial activity maps slightly improved the median PF correlation, PV correlation, and PF shift by 0.072 ± 0.036 (mean ± s.d.), 0.066 ± 0.032 (mean ± s.d.), and 3.35 ± 6.20 mm (mean ± s.d.), respectively. The improvement in PF correlation is significantly higher than would be expected by random chance (p < 10^−5^, 2/4 fish, p < 0.01, 1/4 fish, n.s., 1/4 fish, **Extended Data Fig. 10, Methods**), suggesting a weakly coherent rotation of the place fields. However, even after applying the optimal rotation and nonrigid transformation, PF correlation, PV correlation, and PF shift continues to be significantly different than controls (**Extended Data Fig. 6e**, p < 10^−5^, 4/4 fish for PF correlation and PV correlation; p < 10^−5^, 2/4 fish, p< 0.001, 1/4 fish, n.s., 1/4 fish for PF shift; per-fish one-sided Wilcoxon signed-rank test), suggesting that coherence between place cells is only weakly maintained during remapping.

## Discussion

Place cells, a key computational unit of spatial cognition in mammals, have yet to be convincingly identified outside of mammalian and avian species. The lack of prior evidence for place cells in vertebrate species such as teleost fish, which diverged from mammals ∼450 million ago, suggests one of several possibilities. One possibility is that spatial cognition was subserved by a fundamentally different neural computational network in early vertebrate evolution^19^. In this view, the neural architecture necessary to generate place cells may have evolved sometime between 150 to 400 million years ago, between the emergence of land vertebrates and birds^1,17^. An alternative possibility is that hippocampal and para-hippocampal structures exist in the teleost brain but have not been clearly identified^22^. Indeed, significant controversy exists over the correspondence of the teleost telencephalon to regions in the mammalian forebrain. It has been variously hypothesized that the teleost telencephalon mostly corresponds to a primitive limbic system, a para-hippocampal system, or a primordial neocortex^21,22^. Without a clearly defined location for a hippocampal homolog in the teleost telencephalon, sparse electrophysiological recordings of neural activity in the telencephalon may have limited the discovery of a comprehensive network of place-encoding neurons.

In this study, by comprehensively recording neurons across the larval zebrafish brain in freely swimming animals, we identify place-encoding neurons that are significantly enriched in the zebrafish telencephalon. These neurons collectively encode the animal’s position in space, and their place fields appear to be determined by both idiothetic and allothetic information. Similar to mammalian place cells^8,45,46,51^, without interrupting path integration, place fields in the zebrafish telencephalon remain stable despite significant changes in visual illumination and removal of landmarks. Conversely, after interruption of path integration by removing the animal from the arena, we find that even partially maintained allothetic information (e.g., landmarks or geometric shape) is sufficient to enable recovery of the spatial map. In contrast, complete disruption of path integration, landmarks, and geometric shape lead to significantly greater remapping.

Collectively, the evidence points to multiple sources of input (both idiothetic and allothetic) that stabilize the activity of place-encoding neurons in the teleost telencephalon, suggesting that the multimodal integration properties of place cells may be evolutionarily conserved from early vertebrates to mammals. Confirming the existence of such neurons in the teleost telencephalon fills a long-standing gap in the conceptual framework for spatial cognition in nonmammalian species, and substantially changes the models of spatial cognition that are plausible for primitive vertebrate brains.

In addition to place cells, our data suggests that the teleost telencephalon also contains a small number of border vector cells. Border vector cells have recently been described in goldfish and suggested to be the primary component of spatial computation in lieu of place cells in fish^19^. However, we find that only 1.60 % of spatially selective neurons in the telencephalon fit the definition of a border vector cell^10,11^. Thus, our data suggest that while BVCs exist in the telencephalon, they are distinct from more spatially selective neurons in the telencephalon.

This work sets the stage for an exciting series of questions going forward. First, our characterization of place cell properties so far has focused on early exposure to an initially novel environment. In mammals, it has been demonstrated that repeated exposure to a spatial arena over the course of several weeks leads to 1) stabilization of the place code, 2) greater spatial discrimination between arenas with distinct geometries, and 3) prioritization of allothetic over idiothetic information. Future extensions to tracking microscopy that enable imaging of juvenile zebrafish after weeks of prior experience would be helpful in determining whether these experience-dependent changes in the place code are evolutionarily conserved in teleosts.

Second, not all remapping experiments are currently feasible in larval zebrafish due to the limitations of existing imaging technology. Multisession remapping experiments across days and across multiple chambers are currently limited by the photobleaching rate of fluorescence imaging and will require more light-efficient microscope designs. Replicating mammalian remapping experiments that take place in distinct recording rooms will require the construction of multiple tracking microscopes.

Third, while we find that the place code is stably maintained upon a significant drop in environmental illumination (from 7.05 µW/mm^2^ to < 0.01 µW/mm^2^), we cannot eliminate a small blue light (0.5 mm in radius) locked to the center of the fish brain for neural imaging. We estimate the scattered blue light to be 0.14 µW/mm^2^ within 5 mm from the center of the fish brain. To achieve complete darkness, we would need a tracking system with invisible two photon excitation for neural imaging. In the future, with a two-photon tracking microscope, one could potentially measure place field drift in complete darkness.

Fourth, while spatial cognition has been extensively demonstrated in adult teleosts, only a handful of studies have found evidence of learned place preference in larval zebrafish^52,53^. This suggests several possibilities that should be more fully investigated in the future. One possibility is that learned place preference in larvae requires an appropriate unconditioned stimulus (US). The most commonly used US is an electric shock, which can cause seizure-like activity across the larval brain and may disrupt the process of memory formation^53,54^. Indeed, using more naturalistic rewards (e.g., social reward), place preference has been demonstrated in larval zebrafish between 6 and 8 dpf^52^, but additional work will be needed to fully distinguish between spatial and stimulus-response learning. A second possibility is that place cells mature before spatial cognitive abilities come online. This would be consistent with findings in mammals that place cells can be detected in the hippocampus at P15, but allocentric spatial memory becomes apparent only around P21^55^. If the same phenomenon holds in zebrafish, it might suggest a conserved ontogeny of spatial cognition across vertebrate evolution.

Lastly, we find that place fields are not completely uniformly distributed throughout the environment, but some place cells show correlated activity with one another. Recent work in adult rodents has suggested that spatial tuning across the hippocampus is not independent but may form correlated assemblies^35–37,43^. This implies that the hippocampus may draw upon “preconfigured network states”^35^ in the representation of novel environments. Our data in zebrafish may be consistent with this view and suggests that latent assembly structure may have been a feature of spatial computation that emerged early in vertebrate evolution. However, future studies combining functional imaging with the underlying synaptic connectivity of the teleost telencephalon will be needed to further refine this hypothesis.

## ACKNOWLEDGEMENTS

We thank R. Trivedi, S. Aimon, C. Heller, L. de Sardenberg Schmid, D. Cassidy Nolan for discussion and feedback. This work was financially supported by the Max Planck Institute for Biological Cybernetics.

## AUTHOR CONTRIBUTIONS

J.M.L and D.N.R. conceived of the project on identifying cells with spatial information in the larval zebrafish brain. J.M.L., C.Y., and D.N.R designed the behavioral chambers and experiments. J.M.L. and C.Y. collected the primary data. C.Y. conducted the primary analysis related to spatial information, encoding, anatomical location, manifold structure, and remapping, as well as data preprocessing. L.M. developed the direct basis decoder, contributed to the design of the morph chamber, and analysis of manifold structure. B.K. contributed to data collection, data preprocessing and analysis for remapping experiments and lights off experiments. M.L. performed the preliminary analysis and first identification of place-encoding cells using data from Marques et al. 2020. D.N.R. designed the software and hardware for all imaging systems. C.Y., L.M., and B.K. contributed to all aspects of the data analysis and manuscript preparation with guidance from J.M.L and D.N.R.

## ONLINE METHODS

### Experimental Setup

#### Simultaneous behavioral and neural imaging by tracking microscopy

Tracking microscopy was performed as described in our previous study^1^. Motion cancellation was performed by a custom 3 axis motorized system, to be described elsewhere (Mohan, Li, and Robson, in preparation). To enable animal tracking, the behavioral chamber is illuminated by four custom light strips consisting of narrow angle 850 nm infrared LEDs (SFH4655-Z, Osram) that deliver infrared light into the chamber by total internal reflection. Ambient white light illumination in the behavioral chamber is provided by an array of wide angle white LEDs (GW PSLM31.FM).

To record neural activity, we used DIFF microscopy to image 1013 x 764 x 150 µm of the brain at cellular resolution at a volume rate of 2 Hz and a frame rate of 200 Hz as described in our previous study^1^. The performance and characteristics of the imaging system have been characterized in depth in our previous study^1^.

#### Chamber construction and experiment design

Several constructions for the chamber wall were used depending on the complexity of the experiment. The top and bottom of all behavioral chambers are made of glass to allow optical access from above and below^1^. Between the glass plates, 1 mm thick gas-permeable polydimethylsiloxane (PDMS; Sylgard 184, Dow Corning) walls were used to create a water-sealed rigid body as described previously^1^. PDMS chambers were cut using a computer-controlled blade and finished by manual cuts. The chamber was filled with E3 water before fish loading (see next section on **Fish loading, removal, and reloading**). After fish loading, we gently closed the chamber by sliding on a top cover slip. Excess fluid was removed to create a water-tight seal between the PDMS and the two glass surfaces. Each imaging session lasted for 45-100 min to ensure sufficient spatial coverage of the arena (each experiment is described in more detail below). We do not include the fish in our analysis if it is quiescent for more than 20 minutes during the experiment. If not specified explicitly, all the chambers are initially novel to the fish.

For whole chamber rotation experiments (**Fig. 4a**) and rectangle to circle experiments (**Fig. 5m**), the outer wall (1 mm thick) is made of translucent white PDMS, cut into the desired chamber shape. Landmarks are constructed from black PDMS pieces embedded into the outer wall. A clear PDMS inner wall (0.5 mm thick, 1.5 mm wide) placed alongside the outer wall prevents the fish from directly interacting with the outer wall or landmarks. For the chamber rotation experiments, animals were first imaged for 90 min in the rectangular chamber (session 1), then the entire chamber was slowly rotated over 10-20 s as a rigid body without removing the animal. Animals were then imaged for another 60 min (session 2).

For experiments that required rotation of the chamber wall without removing the animal or disassembling the chamber (**Fig. 4f**), an outer PDMS wall with a center square cutout of 60 × 60 mm was used only for water-sealing the chamber. An inner chamber wall (of distinct size and shape from the outer PDMS wall) was constructed out of laser-etched plastic with embedded stainless-steel pieces (grade 430), which enables remote repositioning of the plastic chamber wall using small magnets (3-5 mm in diameter) below the bottom glass of the chamber. Landmarks (when used) were acrylic-painted onto the wall of the plastic chamber. The plastic chamber wall was placed inside the square cutout of the outer PDMS wall. Animal movement is restricted to within the plastic chamber walls. Given the flexibility of this chamber design, we also used it for experiments in **Fig. 5a, e, i**.

In the wall rotation experiments, we first recorded the animal for 45-90 mins (session 1), then the entire chamber assembly was removed from the microscope. Using small magnets underneath the bottom glass of the chamber, we then carefully rotated the chamber walls by 180° before replacing the chamber assembly on the microscope. During the manipulation, we ensure that the walls do not make strong physical contact with the fish. However, due to manual handling and unpredictable fish movement, it is possible that there is a gentle contact without shifting the fish position. Post wall rotation, the animal was recorded for another 45-75 min (session 2).

For wall morphing experiments (**Fig. 4o**), we constructed a morph chamber consisting of 8 pieces of stainless steel (12 × 2 × 1 mm) linked by a flexible ring of silicone (1 mm wide and 0.5 mm high). The stainless-steel pieces can be remotely repositioned using small magnets below the bottom glass of the chamber (as described above), allowing flexible and gradual morphing of the chamber wall. As described above, an outer PDMS wall with a center square cutout of 60 × 60 mm was used only for water-sealing the chamber, and the morph chamber was placed inside the square cutout. Animal movement is restricted to within the morph chamber. We first arrange the morph chamber into a radially near-symmetric octagon and recorded the animal for 45-65 min (session 1). Using small magnets underneath the bottom glass of the chamber, we then morphed the chamber wall into an ellipse. During the manipulation, we ensure that the walls do not make strong physical contact with the fish. Post morphing, the animal was recorded for another 45-70 min (session 2).

For border insertion and removal experiments (**Extended Data Fig. 3b**), we used a PDMS chamber with a 84 × 15 mm center cutout in which the animal is free to move. At the midpoint of the long axis of the chamber, we created a hidden rectangular pocket (21.5 × 2 mm) containing a stainless-steel rectangular bar (20 × 1.5 × 0.5 mm). Using a small magnet, the stainless-steel bar can either be inserted into the chamber (creating an extra border wall) or hidden from the chamber. In session 1 (45-95 min), the movable wall is hidden from the animal. After session 1, the movable wall was inserted into the interior of the chamber, and the animal was recorded for 45-95 min (session 2). After session 2, the movable wall was again retracted and hidden from the animal, and the animal was recorded again for 45-90 min (session 3).

For the landmark removal experiment (**Fig. 4k**), we used a PDMS chamber with a 25 × 25 mm center cutout. On one edge of the chamber, we created two additional PDMS pockets (100 × 2.5 mm) that each contained a 70 × 2 × 0.5 mm white (acrylic painted) stainless-steel rectangular bar. Landmarks (vertical black stripes) are painted on a portion of one of the stainless-steel bars (**Fig. 4k**), while the opposing stainless bar is completely white. In session 1 (75-90 min), landmarks were positioned to be visible to the fish. After session 1, by applying small magnets to the bottom glass of the chamber, the stainless-steel piece with landmarks was remotely moved into the PDMS pocket, effectively hiding the landmarks from the animal within the chamber. The opposing stainless-steel bar was not moved. The animal was then recorded again for 60-100 min (session 2).

#### Fish loading, removal, and reloading

For all experiments, fish are loaded into the chamber using a glass or plastic pipette. For experiments with fish removal between the first and second sessions (**Fig. 5**), we used the following procedure. After the first 60-90 min recording (session 1), the chamber was removed from the microscope. We slowly inject some E3 water underneath the top cover glass to loosen the bond between the top cover glass and the PDMS walls. This enabled us to smoothly slide away the top cover glass and remove the fish. In between sessions, the chamber was thoroughly cleaned with soap and isopropanol, and then pressure-rinsed for 2-5 min. For two fish (**Extended Data Fig. 6g**), we rotated the bottom glass relative to the chamber walls without cleaning to scramble any potential olfactory cues. We reloaded the fish into the chamber, at a different heading and position than the heading and position at which the animal was removed. Post fish loading, the chamber was again sealed and transferred to the microscope for a second imaging session (45-100 min).

#### Lights on to lights off experiments

To study how the place fields are influenced by the luminance of the environment, we designed a light-dark protocol. After 60 min of recording the fish exploring a circular chamber made of PDMS, we turned off the white LEDs providing visible illumination to the chamber. Turning the white LEDs off results in a luminance change from 7.05 µW/mm^2^ to < 0.01 µW/mm^2^, measured at the center of the chamber. To ensure that the transition to lights off did not generate an aversive response, the luminance was changed gradually over 1 min. Recording was then resumed for an additional 60 min.

We note that a small blue spotlight with a radius of 0.5 mm is always centered on the brain of the fish in all conditions, due to its necessity for neural imaging. We measured the light scattering from this spotlight to be 0.14 µW/mm^2^ within 5 mm of the source.

Light intensity measurements were performed using a photodiode-based optical power sensor with a known detector area (S130C, Thorlabs).

### Image registration

#### High resolution offline registration of fluorescent brain volumes for each animal

Each fluorescent image from the tracking microscope was registered to a high-resolution reference brain volume collected from the same animal. An in-depth description and characterization of the registration pipeline was published in our previous study^1^. Briefly, an initial coarse registration is obtained by optimizing a 3-D rigid transformation that maps the moving image to a (possibly tilted) plane within the reference brain volume. This planar surface is then finely subdivided into a deformable surface that is locally adjusted within the reference volume using a regularized piecewise affine transform.

#### Registration to a common reference brain across animals

We use the Computational Morphometry Toolkit (CMTK)^2^ to register each animal to a common reference fish. An atlas fish^3^ was selected to serve as the common reference brain. Each brain was registered onto the reference brain in a series of steps: initialization, rigid, full affine, warp. The coordinate transformation for each individual cell was then saved. The command line commands are listed below:

*cmtk make_initial_affine --centers-of-mass moving_image fixed_image initial.list*

*cmtk registration --initial initial.list --nmi --dofs 6 --dofs 12 --nmi --exploration 8 --accuracy 0.8 -o affine.list moving_image fixed_image*

*cmtk warp --nmi --threads 160 --jacobian-weight 0 --fast -e 18 --grid-spacing 100 --energy-weight 1e-1 -- refine 4 --coarsest 10 --ic-weight 0 --output-intermediate --accuracy 0.5 -o warp.list affine.list*

*cmtk reformatx --pad-out 0 -o out_image --floating fixed_image moving_image warp.list*

*cmtk streamxform warp.list <cell_coordinates.txt > cell_coordinates_registered.txt*

### Image analysis

#### Extraction of neural activity by Non-negative Matrix Factorization

Non-negative Matrix Factorization (NMF) separates each cell into two components – a spatial footprint and a time varying activity component, as described in our previous study^4^. Briefly, after registration, we applied constrained Non-negative Matrix Factorization^5^ to our whole brain datasets with nuclear localized fluorescence. NMF was performed for each axial section of a given brain volume. Our axial sections of the reference volume are separated by 2 µm, and the average diameter of a zebrafish cell is approximately 5 µm. Thus, there can be double counting of centroids if a cell spans more than one axial plane. Cell centroids were detected throughout the entire reference brain volume, but were only included in downstream analysis if they were sampled in at least 30% of timepoints throughout the imaging session. We merged these cell centroids belonging to the same cell based on close spatial proximity (horizontal distance ≤ 1.4 µm, vertical distance ≤ 2 µm) and highly correlated activity (> 0.7). After merging, the number of cells is 73,621 ± 10,558 (mean ± s.d.), decreasing by 28.4% ± 2.6% (mean ± s.d.) across animals.

The fluorescence baseline for each merged cell was estimated using the 10^th^ percentile within a 30 min sliding window. Baseline-corrected fluorescent traces Δ*F*(*t*) were then obtained by subtracting the estimated baseline from the raw fluorescent traces.

### Spatial information analysis

#### Spatial activity map

A spatial activity map was generated for each neuron, representing the mean neural activity at each spatial location. To calculate the spatial activity map, the chamber was divided into square bins (side length = 1.2 mm), then the summed neural activity and occupancy time were calculated for each bin. This resulted in two matrices: a summed neural activity matrix and an occupancy matrix (matrix entries correspond to spatial bins). We then applied a boundary-constrained Gaussian filter (standard deviation = 1 bin, and the boundary was defined by the chamber boundary) to these two matrices. The place field map was calculated by dividing the filtered summed neural activity matrix with filtered occupancy matrix to obtain the filtered average activity in each spatial bin. When the fish was stationary (speed < 0.1 mm/s), the corresponding frames were not included in the calculation of the place field map. For all experimental spatial activity maps (e.g. for comparison of spatial maps across sessions), we exclude the first 15 min after initial exposure to the environment in session 1. For the within-session control (comparison between the first and second half of session 1), we separately generated spatial activity maps for the first half and second half of session 1. The first 15 min were not excluded in the within-session control to ensure enough coverage of the environment by the fish trajectory.

#### Spatial information and specificity

From the derived place field maps, spatial information can be used to quantify how much information a cell has about the location of the animal, in units of bits per second^6^. For each cell, spatial information *I* was calculated as

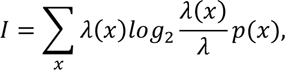

where *x* is a spatial bin, *p*(*x*) is the probability that the fish is in spatial bin *x*, *λ*(*x*) is the mean activity of the cell when the fish is in spatial bin *x*, and *λ* is the average neural activity, computed as *λ* = ∑_*x*_ *λ*(*x*)*p*(*x*).

Based on the equation above, cells with high average neural activity tend to have higher spatial information. To normalize for this, we calculate the specificity *s* as

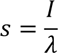

In other words, spatial information divided by average neural activity, resulting in units of bits per activity unit (a.u.).

Due to our baseline correction, bins occasionally have negative average activities. Such bins as well as those with less than 1 s total occupancy time after the Gaussian filter were not included in the calculation of spatial information and specificity.

#### Identification of place fields

The firing field (or place field) of each neuron was defined as the set of spatial bins with activity above 80% of the peak activity (95^th^ percentile) of the spatial activity map. The location of the place field is represented by its center of mass (COM) for cells with a single place field. For cells with multiple place fields (as distinct components in the map), we use the COM of the component (>20 bins, to avoid spurious place fields due to noise) with the highest peak activity (defined as 95^th^ percentile of the component) as the location of its place field. Only place cells with a place field size (after excluding components ≤ 20 bins) of less than 30% of the chamber size are included in maps of the distribution of place fields and in the analysis of place field shift.

#### Rigid and nonrigid registration of spatial activity maps across sessions

Experiments were conducted with various chambers, sometimes in varying orientations, sometimes before and after morphing the chamber into different shapes. To compare spatial activity maps across these chambers, we developed methods to register maps across sessions. When there was no change in the chamber wall geometry (e.g. whole chamber rotation), registration was performed by rotating and translating the session 2 activity map (i.e. a rigid transformation) so that activity in both sessions was represented in terms of the spatial bins of session 1, thus facilitating comparisons (see **PV correlation, PF correlation and PF shift** below).

Otherwise, we performed a nonrigid transformation to register the spatial activity maps across sessions. Our strategy was first to establish a correspondence between the chamber walls of both sessions, then to map each spatial bin of session 2 to a set of spatial bins in session 1, and finally to represent the activity of session 2 in terms of the spatial bins of session 1. Each of these steps is described in more detail below.

First, we detected the walls of the chambers in both sessions and defined a set of anchor points on the chamber walls. Since the precise number of detected wall points could differ between sessions, we used linear interpolation to upsample the wall points of the session with less points, such that the number of anchor points was the same across sessions. The correspondence between anchor points across sessions was established either based on landmarks (e.g. the wall morphing experiment with fish removal, **Fig. 5i**), or based on the polar angle of the wall within the microscope reference frame in experiments with no clear match between geometry or landmarks (e.g. wall morphing without animal removal, **Fig. 4o**, and the rectangle to circle experiment, **Fig. 5m**), or by systematically considering every rotation of the anchor points in session 2 relative to session 1 (**Extended Data Fig. 6c-d, Extended Data Fig. 10**, see next section on **Nonrigid transformation with the best rotation angle**).

Next, for each spatial bin center (*x*, *y*) in session 2, we compute a corresponding location (*x*′, *y*′) in session 1 using a center of mass procedure. Specifically, for each anchor point (*x*_*a*_, *y*_*a*_) in session 2, we associate a weight 1/*d*^2^, where *d* is the distance between the anchor point (*x*_*a*_, *y*_*a*_) and the spatial bin center (*x*, *y*) in session 2. We then use the previously established wall correspondence between both sessions to transfer these weights to the anchor points in session 1. The corresponding location (*x*′, *y*′) in session 1 is then defined as the center of mass of the session 1 anchor points (i.e. the weighted sum of the session 1 anchor points, divided by the sum of the weights).

Finally, we represent the activity of session 2 in terms of the spatial bins of session 1. The computed location (*x*′, *y*′) in session 1 is generally in between the spatial bin centers of the session 1 activity map, so we identify the 4 × 4 spatial bins with bin centers (*x*, *y*) with *floor*(*x*^′^) − 1 ≤ *x* ≤ *ceil*(*x*^′^) + 1 and *floor*(*y*) − 1 ≤ *y* ≤ *ceil*(*y*′) + 1. In this way, we associate each session 2 spatial bin with 4 × 4 spatial bins in session 1. This procedure ensures that each session 1 spatial bin is associated with at least one session 2 bin. We then average the activity of all session 2 bins that are associated with a given spatial bin in session 1, yielding a representation of the activity of session 2 in terms of the spatial bins of session 1.

#### Nonrigid transformation with the best rotation angle

Nonrigid transformation of the session 2 maps described above are generally applied assuming a rotation angle of 0 between sessions (e.g. **Fig. 4r and Fig. 5l, p**). To systematically investigate whether a coherent map rotation has occurred between sessions (**Extended Data Fig. 6d**), 72 incremental rotations (covering 360°) were applied to the session 2 anchor points, followed by nonrigid registration as described above. The best rotation angle is identified by the maximum PF correlation (see **PV correlation, PF correlation and PF shift**) across sessions. We refer to this procedure as “nonrigid transformation with the best rotation angle”. Note that this is also done for the early-late control in **Extended Data Fig. 6c, e**, and **Extended Data Fig. 10**.

To test whether the improvement in PF correlation from the “nonrigid registration” assuming an angle of 0 to “nonrigid transformation with the best rotation angle” is significant, we shuffle the cell identity in session 2. Post shuffle, we then compare the improvement in PF correlation from the “nonrigid registration” assuming an angle of 0 to “nonrigid transformation with the best rotation angle”. This was repeated 1000 times to generate a null distribution. Then the *p*-value in **Extended Data Fig. 10** is calculated by counting the proportion of times the real data improves by more than the shuffle control.

#### PV correlation, PF correlation and PF shift

To compare the population level activity across sessions, a population vector correlation (i.e. PV correlation) was computed for each spatial bin that is shared across sessions. 1) For each neuron, we first compute a Δ*F*/*F* spatial activity map for each session. To do this, we estimate a baseline fluorescent signal for each map by taking the mean of the spatial bins with fluorescent signal below 20^th^ percentile. Then the Δ*F*/*F* of each spatial bin is computed as follows:

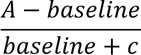

*A* is the mean fluorescent signal for a spatial bin. A pseudocount *c* is added when the baseline is below 10. For each spatial bin, we obtained two vectors of population level activity (one for each session). The length of the vector is the power of the union of the telencephalic place cells identified from either session. Correlation between the two activity vectors yields the PV correlation for a given spatial bin.

To determine the similarity of spatial activity maps, we computed the correlation between the spatial activity maps (i.e. PF correlation or spatial correlation). Correlation was performed on the spatial bins that are shared across both maps and normalized by the mean and variance of each map.

To measure how much place fields shift across sessions, for each neuron, we identified the COM of its place field in each session. We define PF shift as the distance between the COM of the cell’s place field in session 1 and session 2. Only cells with confined place fields (firing field size less than 30% of the chamber size, see **Identification of place fields**) in both sessions are included for this analysis.

For the within-session control, we generate two separate spatial activity maps, corresponding to the early and late half of session 1. The PV correlation, PF correlation, and PF shift are computed from the within-session control spatial activity maps with the same procedure as described above. For the comparison to the control, we make sure that the same set of neurons (for PV correlation, PF correlation, and PF shift) and spatial bins (for both PV correlation and PF correlation) are used. One-sided Wilcoxon signed-rank tests are used to measure whether PF correlation and PV correlation are significantly smaller than the control, and whether PF shift is significantly larger than the control.

##### Specificity z-score and population specificity z-score

To test the significance of the specificity of each cell, we use circular permutation to construct a null distribution. For each cell, we define the set of valid timepoints as the frames where neural activity was recorded and the fish movement speed is > 0.1 mm/s. Then a null distribution for specificity is estimated by measuring the specificity after circularly permuting the neural activity vector within the valid timepoints by 1000 offsets (each offset is 0.5 s, so it covers from −250 s to 250 s). The specificity of a given cell is converted to a specificity z-score by subtracting the mean specificity of the null distribution and dividing by the standard deviation of the null distribution. To test the significance of the specificity of a cell at the population level, a population specificity z-score is also calculated by subtracting the mean specificity of all recorded cells and dividing by the standard deviation of the specificity of all recorded cells. To be classified as a place-encoding cell, a cell is required to have a specificity z-score larger than 5, a population specificity z-score larger than 3, and a specificity value of > 0.01 bits/activity unit. For the three criteria we use (specificity z-score > 5, population specificity z-score > 3, specificity > 0.01 bits/activity unit). The percentage of cells that pass each criterion are: 1.88 ± 0.22 % for (population specificity z-score > 3), 17.71 ± 7.32 % for (specificity z-score > 5), and 87.29 ± 7.17 % for (specificity > 0.01 bits/activity unit).

#### Blurring of the spatial activity map

Blurring is the broadening of a place field at a given point on its boundary in the radially outwards direction. To estimate this blurring effect of place fields due to the speed of the fish combined with the calcium indicator kinematics, we compute an upper and lower bound. The lower bound is computed by assuming a 10 Hz firing rate and a speed of mean minus standard deviation. The upper bound assumes a firing rate of 100 Hz with a speed of the mean plus the standard deviation. The speed distribution consists of pooled speed data from 7 fish (**Extended Data Fig. 1f**). The data for the half-decay times for different firing rates are taken from Chen et al., 2013^7^. The half-time of the temporal fluorescence decay is multiplied by the speed value to get the half-distance for a spatial fluorescence decay (**Extended Data Fig. 1g**). Of this exponential spatial decay, the distance needed for a decay down to 80% of the starting value is computed (**Extended Data Fig. 1h**). This matches our definition of place field and is used as blurring estimate.

#### Isomap and quantification

Isomap^8^ embedding was performed using scikit-learn^9^ with n_neighbors = 100. A rectangular matrix representing the population activity of all telencephalic place cells in a span of 30 min was constructed, then an isomap manifold was fit to the data (with each point in the manifold representing the population activity at one timepoint), and finally a 2D embedding was extracted for visualization in **Fig. 3a** and computation in **Fig. 3b**. Since the implementation does not handle missing data, we filled in missing values in the activity trace for individual cells with the nearest preceding available value. For **Fig. 3**, the early (0-30 min) and late (60-90 min) windows of population activity data were fit separately and therefore have different 2D embeddings. For **Supplementary Video 2** we fit an isomap manifold to the last 30 min of imaging session 1 (before chamber rotation) to establish stable axes for 2D embedding and then repeatedly transformed each window of 30 min of population activity data into this embedding. Standard exclusion criteria based on movement were applied (see section on **Spatial activity map**).

To quantify the relationship between the 2D manifold and the physical position of the fish, we use neighbor distance, which quantifies the degree to which local neighbors in the manifold space are also local neighbors in the physical space of the chamber. In a given 30 min time window, we analyze each timepoint in the 2D manifold space (the “seed point”) by selecting its 30 nearest neighbors in the manifold space, then measuring the physical distance in the chamber between those 30 points and the seed point, and averaging to obtain the mean physical neighbor distance of the seed point. We then compute the overall average physical neighbor distance by averaging the physical neighbor distance across all timepoints in the manifold. As a baseline, for each seed point, 30 random neighbors in the manifold are chosen, and then physical neighbor distance is computed as before.

For analyses comparing between two sessions or between the early and late intervals of a session (**Fig. 3**), the occupancy of each spatial bin was equalized by subsampling. That is, for each spatial bin, we identified the session or time interval having a lesser number of timepoints, and randomly subsampled from the session or time window with more timepoints, so that both time windows had the same number of timepoints in each spatial bin.

#### Direct basis decoder

The direct basis decoder^10^ is a linear decoder with no free parameters that predicts the animal’s position in each individual time point by a linear combination of the place maps, weighted with the corresponding place-encoding cell’s activity. All decoding and map construction was only done on timepoints where the fish was moving (fish speed > 0.1 mm/s). Even though no parameters had to be learned, it is still necessary to compute the place field of each cell. To avoid circularity between computing the place fields and testing the decoder, we used a cross validation scheme in which the neural data was divided into non-overlapping one-minute chunks. To test the decoder on each one-minute chunk, we first computed the place fields without including the test chunk or its two neighboring chunks. The predictions of all one-minute chunks were then concatenated as the prediction of the whole dataset. The decoder error was then computed as the mean distance between the predicted and actual position of the fish across the dataset.

Place fields were constructed as described above using spatial bins (side length = 1.195 mm) and boundary-constrained Gaussian smoothing. Apart from the analysis in **Extended Data Fig. 5c**, the activity used in this map construction is time-shifted by 2 seconds relative to the location of the fish, to counteract possible calcium lag dynamics. All maps were standardized by mean and standard deviation. To prepare the decoder, a 7.5 s boxcar average filter was applied to the activity, and then maps of the most active 30% of cells were weighted by their respective activity and summed to form the decoder map. This nonlinear thresholding step was to cut out low intensity signals that would contribute noise to the decoding. The decoder map was then normalized for density differences in representation. To this end, we calculated a representation histogram across all spatial bins, defined for each spatial bin as the number of place cells that contain that bin in their primary place field (the > 95^th^ percentile component as described above). Since we found that the effect of representation density on decoding error was sublinear, we normalized the decoder map by the third root of the representation histogram. The position estimate for each timepoint was then calculated by the center of mass of the 99^th^ percentile of the decoder map. If not specified differently, 1000 telencephalic cells of highest spatial tuning per animal were used for all decoding analyses. To select the top spatially-tuned cells, we ranked each telencephalic place-encoding cell according to its specificity z-score and also according to its population specificity z-score, and then assigned its final rank according to the worse of the two ranks. This allows for a fair comparison between fish, that may have varying numbers of place cells (**Fig. 1g**). To select the minimum number of cells needed for good decoding, an iterative, greedy algorithm is employed. Starting with zero neurons, we find the single best neuron to add to the decoding set to minimize the resulting decoder error, and then iteratively repeat this same greedy selection procedure to grow the decoding set one neuron at a time.

For the analysis of decoder error by brain region (**Fig. 2e**), the 1000 top spatially-tuned cells were chosen for each defined region (whole brain, telencephalon, mes- and rhombencephalon). The random cell population was randomly selected from cells with a population z-score less than 1, to explicitly not contain any place cells. We defined two baselines for evaluation of the decoder performance. For the uniform random baseline, we measured the average decoder error of a decoder that outputs random positions sampled uniformly from the chamber. This procedure was repeated 1000 times, and the mean decoder error was taken as the uniform random baseline. For the behavior-informed baseline, we measured the average decoder error of a decoder that outputs the center of mass of the fish positions in the chamber.

For all analyses, if not specified differently, only the last 75 minutes of each experiment were used for decoding. In the analysis of the decoder error as a function of number of cells included (**Fig. 2f**), the threshold for spatial tuning was gradually lowered until 10,000 cells were obtained (n = 7 fish). The two fish with less than 10,000 cells in the telencephalon were hence excluded from this analysis. To avoid circularity, for **Fig. 3d**, the shuffled specificity z-score as well as population z-score were calculated for the first and last 30 min of the experiments separately. Decoding for these two time-windows was therefore based on the respective top-ranked place cells within each window.

#### Regression and prediction of neural activity

We used ordinary least squares (OLS) regression to measure how much of the variance in place-encoding cell activity can be explained by different behavioral variables, including physical location, heading and speed. This is a much simpler model compared with previous work using a generalized linear model (GLM)^11^. The behavior variables were discretized into bins, and an indicator function for each bin was added as a regressor in the model. The regression model can be summarized by the following formula:

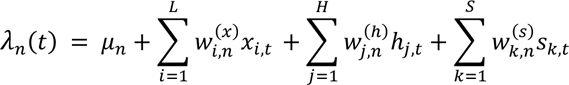

where *λ*_*n*_(*t*) is the baseline corrected neural activity for cell *n* at time *t*, *μ*_*n*_ represents the baseline activity, 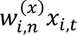 represents the contribution from the physical location of the fish, *x_i,t_* is the activity for spatial bin *i* at time *t*, *L* is the number of spatial bins (15 × 8 = 120), 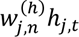 represents the contribution from the heading of the fish, ℎ_*j*,*t*_ is the activity for heading bin *j* at time *t*, *H* is the number of heading bins (24), 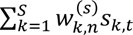 represents the contribution from the speed of the fish, *s*_*k*,*t*_ is the activity for speed bin *k* at time *t*, and *S* = 24 is the number of speed bins. To compare the contributions from spatial location, heading, and speed individually and together, we tested OLS regression in four cases: 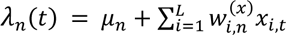, i.e. with only spatial indicator functions, 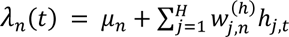, i.e. with only heading indicator functions, 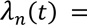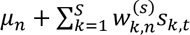, i.e. with only speed indicator functions, and finally the complete equation shown above. All models included a weak L2 regularization on the parameters with a penalty coefficient of 1^−10^. For each regression model, we report the resulting distribution of *R*^2^ values across the telencephalic place cells in **Extended Data Fig. 1b**. Similarly, the distribution of *R*^2^ values across other telencephalic cells is shown in **Extended Data Fig. 1c**. Standard exclusion criteria based on movement speed were applied (see section on **Spatial activity map**).

#### Whole brain maps of place cells

After projecting the location of place cells from all the fish to the same reference fish^3^, we accumulate all the place cells into a 3D histogram representing how many place cells were detected in each 3D bin within the reference volume. Then, we convolve the 3D histogram with a spherical convolution mask with a radius of 5 µm, such that each place-encoding cell contributes an increment of 1 count to all bins within a 5 µm radius. Maximum intensity projections are then computed to visualize the anatomical distribution from multiple views (**Fig. 1f**).

#### Boundary vector cell (BVC) model

To search for cells whose firing properties follow the boundary vector cell model^12^, we fitted an adapted model to all cells in the telencephalon meeting the standard specificity criterion (> 0.01 bits/activity unit for all 3 sessions). The model predicts that the activity of these cells *f* integrates each boundary point (represented in polar coordinates *r* and *θ* relative to the animal) according to

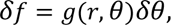

where *g*(*r*, *θ*) represents the firing rate relative to a preferred firing orientation *Φ* and distance *d* from the boundary:

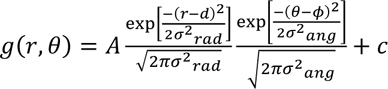

Here, *σ*_*rar*_ *σ*_*rar*_ and *σ*_*ang*_ are the width of radial and angular tuning. Compared with the original model^12^, which treats *σ*_*rar*_ as a variable which linearly depends on *d*, we treat *σ*_*rar*_ as a constant to be fit. *A* is a scaling factor and *c* is the baseline.

For the experiment in which an interior wall was introduced for session 2 and removed for session 3 (**Extended Data Fig. 3b**), we fit the model to the spatial activity map in session 1 and then used the fitted model to predict the map if a wall were inserted in the middle of the chamber. We first identified a set of candidate neurons with the following properties after fitting the model to the session 1 and session 3 maps: a preferred firing orientation *Φ* towards left or right (*Φ* from −45° to 45° or 135° to 225°) in both session 1 and session 3. Given the activity of these candidate neurons in session 1 and 3, the BVC model predicts a duplication of their place fields in session 2. The predicted change in spatial tuning was computed as the difference between the predicted map in session 2 and the fitted map in session 1. We measured the actual change in spatial tuning of each candidate neuron as the difference between the empirically measured spatial activity maps between session 2 and session 1. We then compute the Pearson correlation between the predicted and actual change in spatial tuning, which we refer to as the “prediction performance” of the BVC model for a given cell. Cells with prediction performance > 0.6 are classified as BVCs (**Extended Data Fig. 3d**).

Note that the bin size for spatial activity maps for this experiment is slightly smaller (side length = 1.1 mm) to ensure sufficient resolution to accurately detect and represent the inserted wall. When fitting the model to spatial activity maps, we constrain *d* to be between 0 and 10 bins (assuming BVCs fire close to the boundary), *Φ* between −π and π, both *σσ*_*rraarr*_ and *σ*_*ang*_ between 0 and 10, *A* between 0 and 100, and *c* between −10 and 10 activity units.

#### Neighborhood analysis

To quantify the degree to which potential clustering of place fields is maintained across sessions, we performed a neighborhood analysis for each neuron (**Extended Data Fig. 9**). First, for each neuron, we rank all other neurons by their PF correlation to the neuron of interest in session 1. We define the neurons with the highest PF correlation as “neighbors”, with a systematically varied inclusion threshold of top 2% to top 100%. We define “neighbor retention %” as the number of neurons that remain neighbors in session 2 divided by the number of original neighbors in session 1. The mean “neighbor retention %” across all telencephalic place cells from either session are plotted. This analysis is also done separately for cells whose place field is close to the edge (nearest distance of COM to edge ≤ 3 mm) and cells whose firing field is away from edge (nearest distance to edge > 3 mm). To avoid the contribution from potentially imperfect cell merging, we restricted this analysis to pairs of cells with a minimum anatomical distance > 20 µm. A one-sided Wilcoxon signed-rank test was used to quantify whether “neighbor retention %” in the experiment was significantly worse than a within-session positive control (comparison between the early and late period of session 1). To generate a negative control, we shuffled the cell indices in the second session 1000 times to randomize relationships between neurons. Post shuffle, we calculated the mean “neighbor retention %” across all telencephalic place cells from either session to generate a null distribution (i.e. a shuffle control). Then a *p*-value is calculated by counting the proportion of times the mean “neighbor retention %” is higher than the shuffle control. For these significance tests, we fixed the number of top correlated neighbors to be 10%.

##### Significance tests for sample comparison

Wilcoxon signed-rank tests were used for paired location comparisons with equal sample sizes. Mann–Whitney U tests were used for unpaired location comparisons.

#### Animal care and transgenic lines

Experiments are carried out in accordance with the Animal Welfare Office at the University of Tübingen and the Regierungspräsidium. All experiments in rectangular chambers used Tg(elavl3:H2B-GCaMP6s+/+) with nacre (mitfa-/-) at 6-8 dpf. The rest of the experiments used Tg(elavl3:H2B-GCaMP8s+/+) with nacre (mitfa-/-) at 6-13 dpf. In three of the rectangle to circle experiments, the fish expressed an additional allele of Tg(elavl3:GCaMP6s). Fish were reared on a 14/10 hr light/dark cycle at 25°C. They were maintained in sets of 20 in E3 water and were fed paramecia daily.

#### Computer hardware for data acquisition and analysis

The tracking microscope was controlled by a rack-mounted 12-core workstation with a Geforce RTX 3080 Ti GPU. Offline data analysis was performed using eight rack-mounted 8- to 16-core Linux servers with 4 GPUs each (Geforce RTX 3080 Ti, RTX 2080 Ti, or GTX 1080 Ti).

#### Data availability

The data that support the findings of this study can be made available from the corresponding authors upon request.

#### Code availability

All analysis software was written in Julia 1.7 and CUDA C using the Julia package ecosystem (Optim.jl, PyPlot.jl, Images.jl, Cairo.jl, HDF5.jl, CUDA.jl). GPU processing was implemented for fish tracking, as well as online and offline image registration.

## EXTENDED DATA FIGURES

**Extended Data Figure 1.**
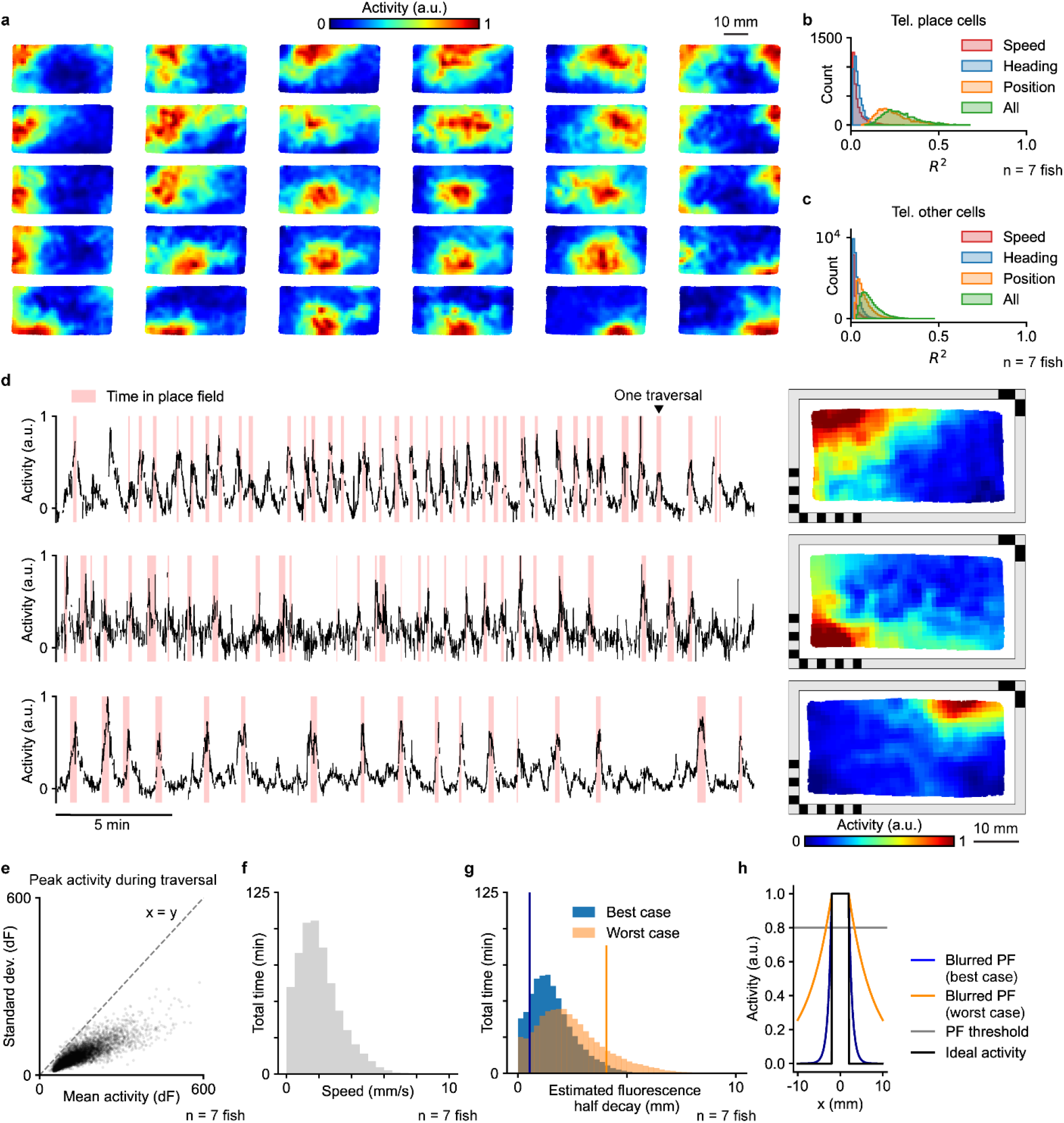
Place cell properties. **a**, Example spatial activity maps at different locations in the chamber. **b**, Modeling the relationship between activity and behavioral variables for telencephalic place cells. *R*² values of individual spatially-tuned telencephalic cells after modeling their activity with ordinary least squares regression (OLS, **Methods**) using only the animal’s speed, heading, or two-dimensional position, as well as using all of them combined (n = 7 animals). **c**, Modeling the relationship between activity and behavioral variables for other cells in the telencephalon. **d**, Activity traces (left) and spatial activity maps (right) for three example place cells. The time that the fish spent inside the place field is marked in red. **e**, Statistics on the firing variability of place cells. One dot represents the mean and standard deviation of the peak activity across all traversals of one place cell. A traversal is defined as a stay inside the place field that is interrupted only by departures shorter than 5 s. Data pooled across 7 animals. **f**, Distribution of swimming speed from the whole experiment from 7 animals, excluding quiescent time periods (swimming speed < 0.1 mm/s). **g**, Distribution of the estimated GCaMP6s fluorescence half decay distance in one linear dimension, based on the data from **f**, and assuming decay half-times of 0.56 s (10 Hz firing rate) and 1.21 s (80 Hz firing rate). Vertical lines indicate the mean - std. dev. for the best case and the mean + std. dev. for the worst case, based on the values of the distributions. **h**, Estimated actual place field blurring in one dimension based on an ideal place field with sharp edges (black line, −2.5 mm to 2.5 mm) and the best and worst cases of half decay distances shown in **g** (blue and orange lines). Place field size is determined by the set of spatial bins with activity above 80% of the peak activity (95^th^ percentile) of the spatial activity map. Blurring to each side is 0.17 mm (best case) and 1.31 mm (worst case).

**Extended Data Figure 2.**
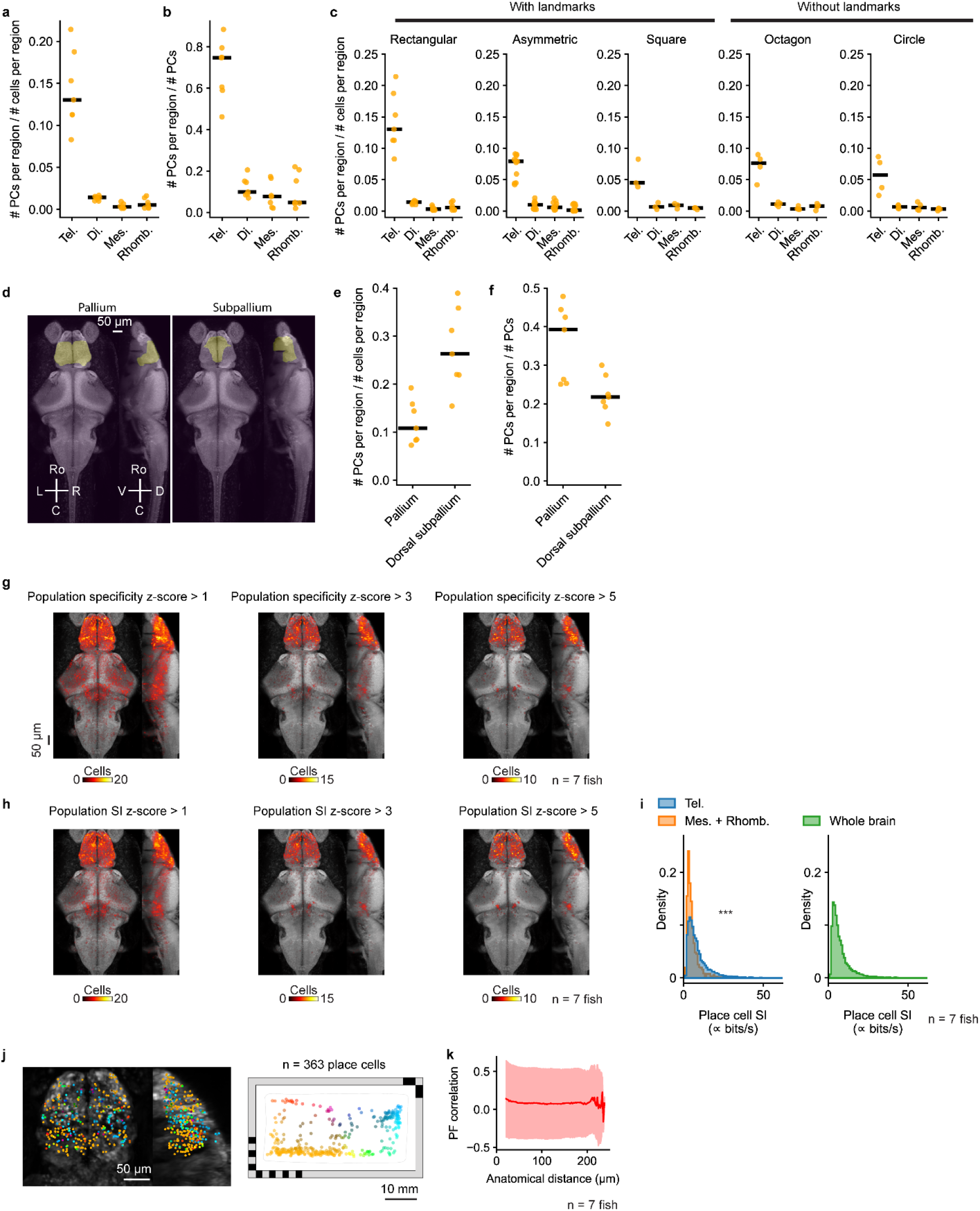
Anatomical analysis of place cells. **a**, Fraction of cells within each brain region that are classified as place cells. Each animal is shown individually (yellow dots, n = 7 animals) along with the median across animals (black line). Tel., telencephalon; Mes., mesencephalon; Di., diencephalon; Rhomb., rhombencephalon. **b.** Fraction of all place cells identified anywhere in the brain that are found in each particular brain region. **c,** Fraction of cells within each brain region that are classified as place cells, shown for five differently shaped behavioral chambers without landmarks: rectangle (**Fig. 1a**), asymmetric shape (**Fig. 5a**), square (**Fig. 4k**), octagon (**Fig. 4o**), and circle (**Fig. 5m**). **d,** Projections of telencephalic subdivisions: pallium and subpallium. Masks imported from the Z-Brain-Atlas. **e,** Fraction of cells in the pallium that are classified as place cells, and fraction of cells in the dorsal subpallium that are classified as place cells. We note that the ventral portion of subpallium extends beyond our imaging volume and is not fully sampled compared with the pallium and dorsal subpallium. **f**, Fraction of all place cells identified anywhere in the brain that are found in the pallium or dorsal subpallium. **g,** Anatomical distribution of cells that would be classified as place cells at different spatial specificity thresholds, based on a population specificity z-score ≥ 1 (left), ≥ 3 (middle), and ≥ 5 (right). Summation of 7 animals overlaid on a reference brain. The black bar indicates 50 μm. **h,** Anatomical distribution of cells that would be classified as place cells at different spatial information (SI) thresholds, based on a population SI z-score ≥ 1 (left), ≥ 3 (middle), and ≥ 5 (right). **i**, Comparison of SI for place cells found in the telencephalon and place cells found in the mesencephalon and rhombencephalon (left, n = 7 animals, p < 10^−5^ in one-sided Mann–Whitney U test), as well as all place cells found in the whole brain (right, n = 7 animals). Since SI was calculated from calcium activity without attempting to convert to spikes/s, the SI values reported here are proportional to classic SI, measured in bits/s, but with an unspecified constant of proportionality. **j**, Anatomical distribution of place cells (left) colored by their place field locations (right). **k**, The relationship between anatomical distance and PF correlation. The solid line indicates the mean. The shaded region indicates the standard deviation. To avoid any possibility of cross talk in the fluorescence between neighboring cells, only cell pairs with anatomical distance > 20 μmare plotted.

**Extended Data Figure 3.**
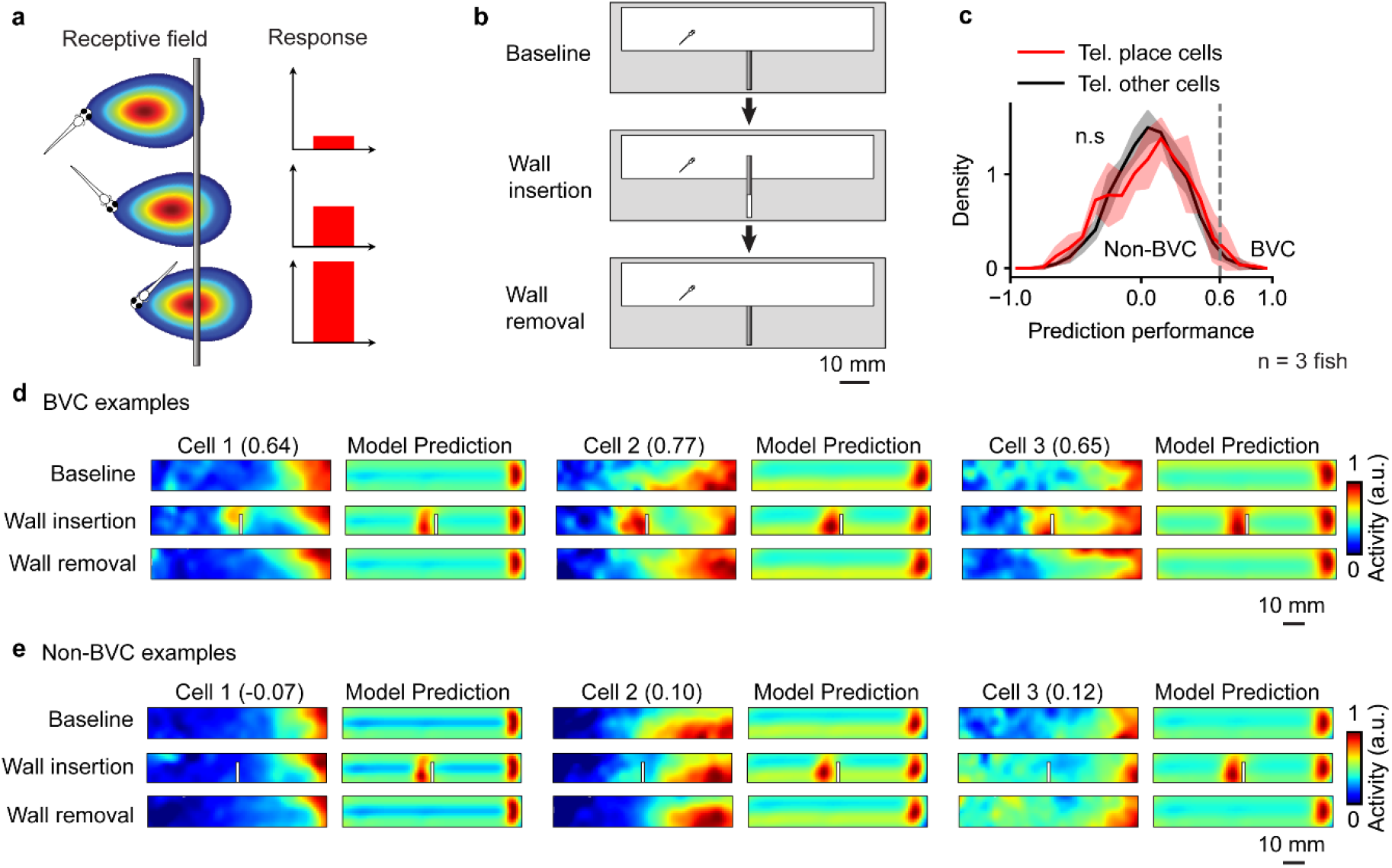
Boundary vector cell analysis. **a**, Schematic of the boundary vector cell (BVC) model. A BVC reaches the highest firing rate when the animal is at its preferred firing orientation and distance to the wall. **b**, Schematic of the experiment to test for BVCs in the larval zebrafish brain (**Methods**). A wall hidden in the middle of the rectangular chamber is inserted between the 1^st^ session (baseline) and the 2^nd^ session of the experiment. Then the wall is removed again for the recording of the 3^rd^ session of the experiment. The BVC model predicts that a BVC with a preferred horizontal firing orientation would duplicate its firing field when a new wall is inserted. **c**, The prediction performance of the BVC model for place cells and other cells in the telencephalon (**Methods**). The prediction performance quantifies how well the BVC model can predict the change in the spatial activity map when the wall is inserted into the chamber. The result of a two-sided Mann–Whitney U test is marked (p ≥ 0.01). The threshold used to identify cells with high BVC model prediction performance is marked (0.60). **d**, Example telencephalic neurons that fit the BVC model well. For each cell, the model is fit to the baseline (1^st^ session) and can be used to predict the spatial activity map for the case of wall insertion or removal (2^nd^ and 3^rd^ sessions). Left: actual spatial activity maps. Right: model-predicted maps. **e,** Example telencephalic neurons that do not fit the BVC model.

**Extended Data Figure 4.**
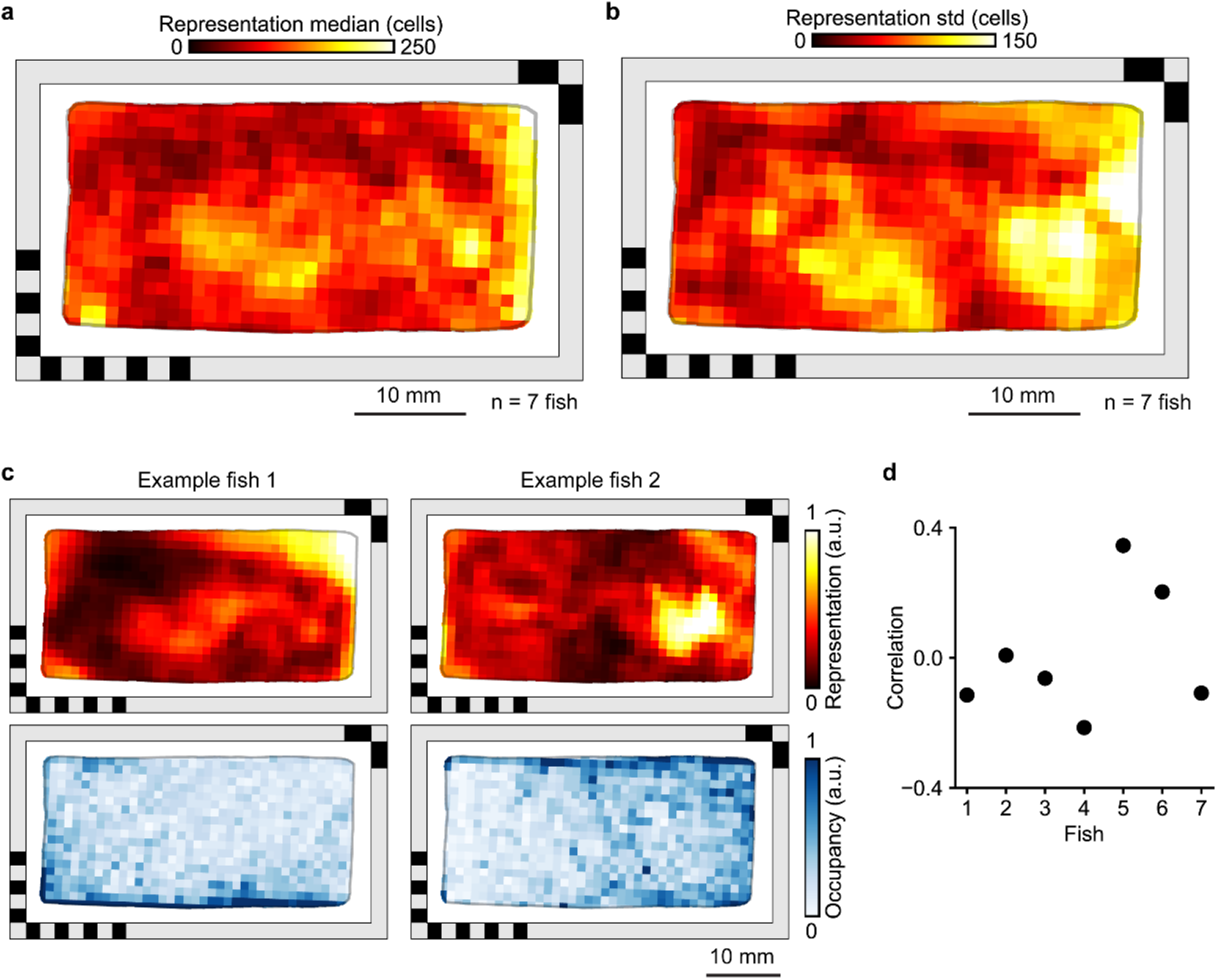
Spatial representation and occupancy. **a**, Distribution of telencephalic place fields encoding different locations in the behavioral arena (same as **Fig. 2a**, n = 7 animals, median across animals is shown). To quantify the density of the place fields across space, each spatial activity map was binarized (**Methods**). **b**, same as **a**, but showing the standard deviation of spatial representation across the 7 animals. **c**, Spatial representation (top) and spatial occupancy (bottom) for two example fish. Representation and occupancy are both normalized. **d**, Correlation between spatial representation and occupancy for 7 different animals.

**Extended Data Figure 5.**
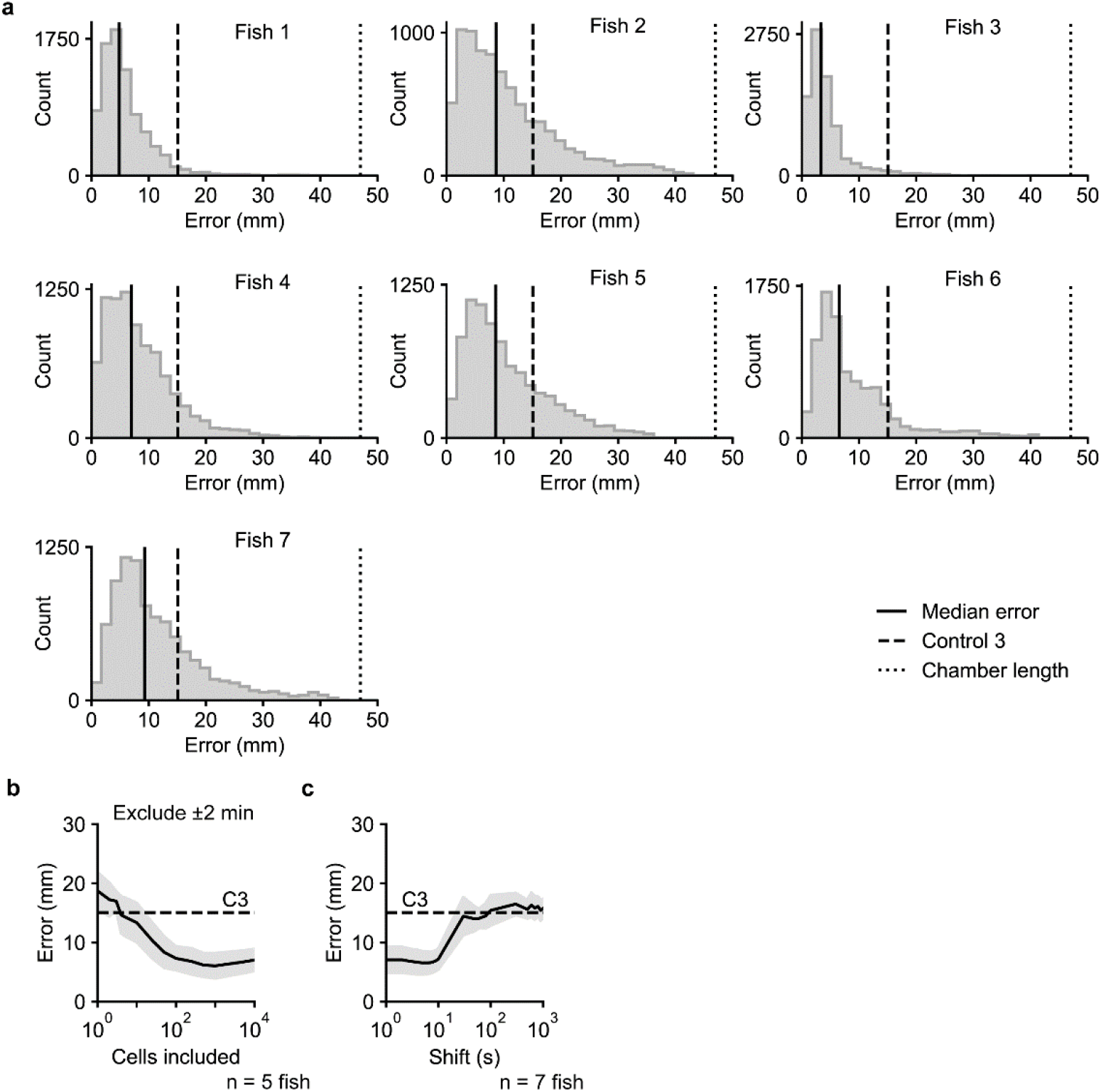
Additional quantification of decoder error. **a**, Distribution of decoder error for 7 animals. Vertical lines indicate the median error (black line), the behavior-informed baseline (15.08 mm, dashed line), and the accessible length of the chamber’s long axis (47 mm, dotted line). **b**, Replication of **Fig. 2f** but excluding from training the 2 min before and 2 min after the 1 min used for decoding (**Methods**). **c**, Decoder error as a function of timing shift between neural activity and animal position (black, mean; gray, std. dev.).

**Extended Data Figure 6.**
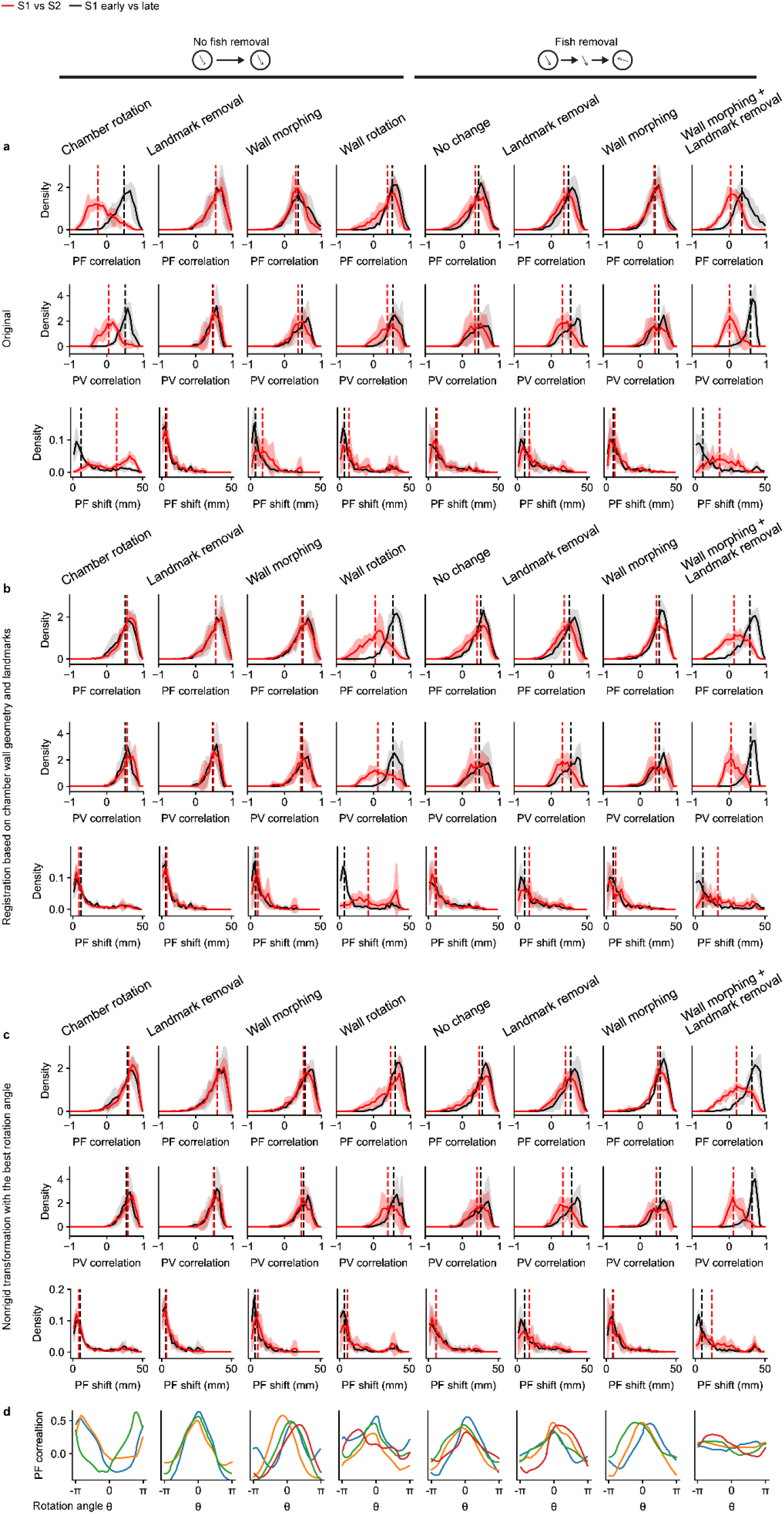
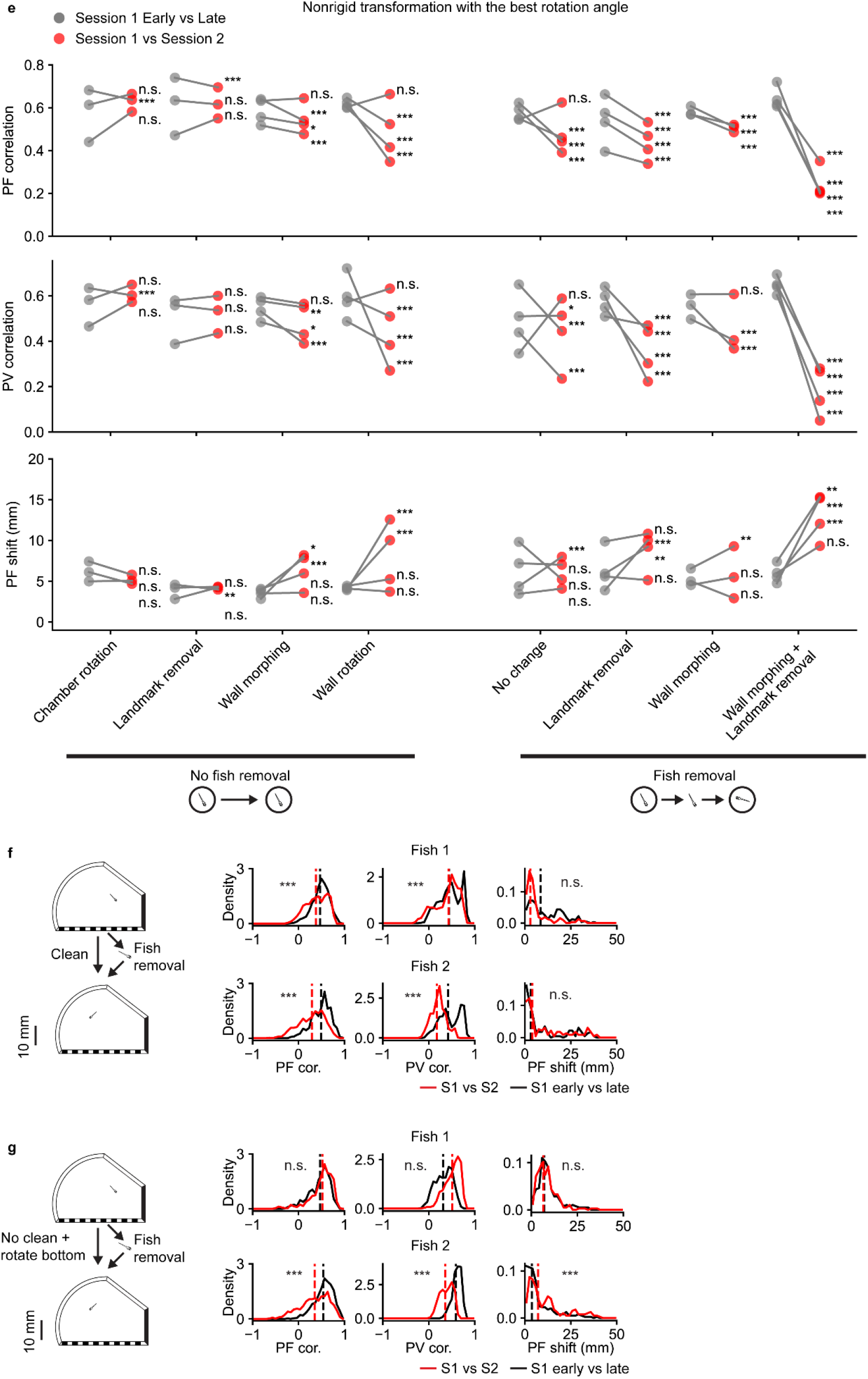
Additional quantification of map comparisons. **a**, Summary of the changes in telencephalic place cells by directly comparing the maps across two sessions in the microscope reference frame. Experiments from **Fig. 4** and **Fig. 5** are represented on the horizontal axis: chamber rotation (**Fig. 4a**), landmark removal (**Fig. 4k**), wall morphing (**Fig. 4o**), wall rotation (**Fig. 4f**), fish removal with no change in the chamber (**Fig. 5a**), landmark removal with fish removal (**Fig. 5e**), wall morphing with fish removal (**Fig. 5i**), and wall morphing with landmark removal and fish removal (**Fig. 5m**). From top to bottom are place field correlation (PF correlation), population vector correlation (PV correlation), and the shift in the place field location (PF shift) (**Methods**). The union of telencephalic place cells from both sessions was used. The comparison between the 1^st^ and the 2^nd^ sessions (red) is plotted against a reference comparison between the early (1^st^ half) and late (2^nd^ half) stages of the 1^st^ session (black). Solid lines are the mean distributions across all fish, and the shaded region indicates the standard deviation across all fish. Vertical dashed lines are the medians of the averaged distributions. **b**, similar to **a** but after registering the maps from the 2^nd^ session to the 1^st^ session with reference to the chamber wall geometry and landmarks (**Methods**). **c**, similar to **b** but the transformation is done by finding the rotation of the reference anchor points that maximizes the mean PF correlation after registration (**b** is equivalent to a rotation angle of 0, **Methods**). **d,** The mean PF correlation as a function of the anchor rotation angle used in the nonrigid transformation, for different experiments (**Methods**). **e**, Summary of changes in the spatial activity maps of telencephalic place cells under different environmental manipulations after applying the nonrigid transformation with the best rotation angle. The median of PF correlation, PV correlation, and PF shift are plotted for each fish. Red dots are comparisons made between the 1^st^ session and the 2^nd^ session. Gray dots are reference comparisons made between the early (1^st^ half) and late (2^nd^ half) stages of the 1^st^ session (also after nonrigid transformation with the best rotation angle). The results of statistical tests are marked for each fish separately (one-sided Wilcoxon signed-rank test, *** for p < 10^−5^, ** for p < 0.001, * for p < 0.01, n.s. for p ≥ 0.01). **f**, Analysis of fish removal experiment as in **Fig. 5d**, but only for individual fish where the olfactory cues were cleaned thoroughly before each recording session with soap and isopropanol. The results of statistical tests for different measures against a control comparison between the early (1^st^ half) and late (2^nd^ half) period of the 1^st^ session are shown for each fish (one-sided Wilcoxon signed-rank test). **g**, Analysis of fish removal experiment as in **Fig. 5d**, but only for individual fish where the bottom was rotated without cleaning to scramble any potential olfactory cues.

**Extended Data Figure 7.**
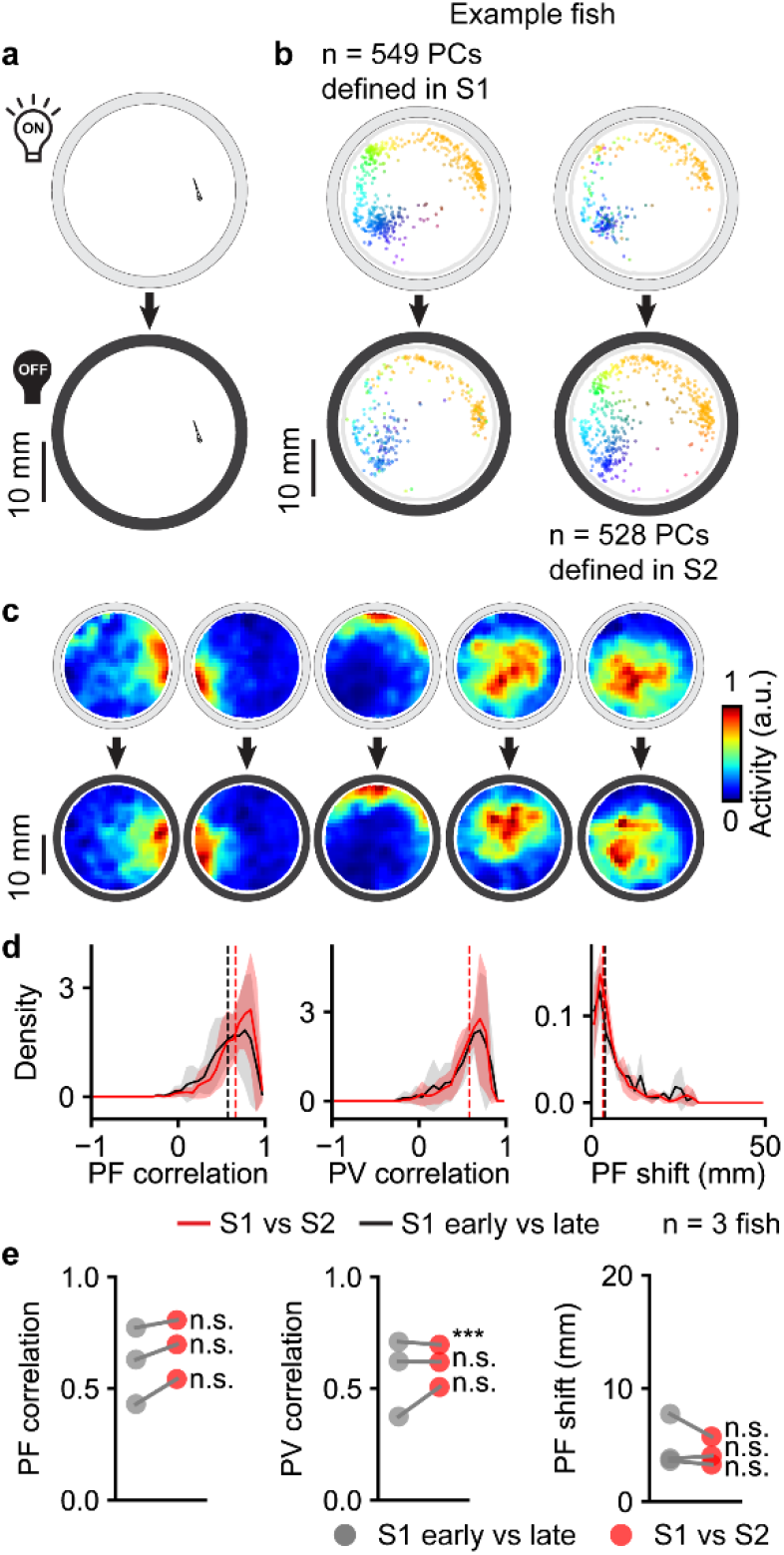
Lights on to lights off experiment. **a.** Schematic of the light-dark experiment in a circular behavioral arena. White light is gradually turned off between session 1 (lights on) and session 2 (lights off, **Methods**). **b-d**, Analysis of light-dark experiment as in **Fig. 4b-d**. **e**, Analysis of light-dark experiment as in **Fig. 6**.

**Extended Data Figure 8.**
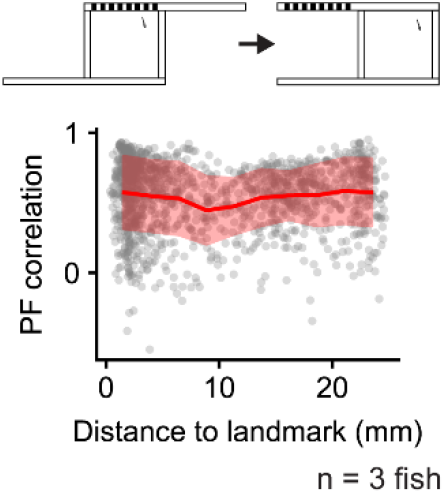
Place field correlation versus environmental features. Top, schematic of landmark removal experiment (**Fig. 4k**). Bottom, PF correlation between recording sessions in the landmark removal experiment as a function of the distance of the place field (represented by its COM) to the landmark.

**Extended Data Figure 9.**
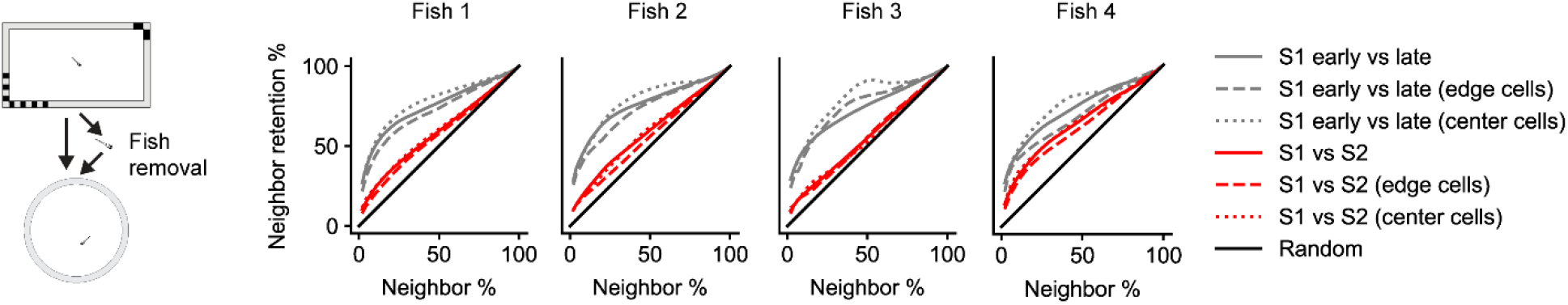
Retention of local neighborhood relationships across environments. For the experiment with both landmark removal and geometric morphing (**Fig. 5m**, schematic shown on the left), we quantify the degree to which neighborhood relationships between place fields are maintained across sessions (“neighbor retention %”). First, for each neuron, we rank all other neurons by their PF correlation to the neuron of interest in session 1. We define the neurons with the highest PF correlation as “neighbors”, with a systematically varied inclusion threshold from the top 2% to the top 100%. We define “neighbor retention %” as the number of neurons that remain neighbors in session 2 divided by the number of original neighbors in session 1. The mean “neighbor retention %” across all telencephalic place cells from either session is plotted for each fish (**Methods**). The comparison between the 1^st^ session and the 2^nd^ session (solid red line) is plotted against a reference comparison between the early and the late stages of the 1^st^ session (solid gray line), as well as a comparison against shuffled data (black solid line). We also subdivide the neurons into 2 groups, depending on the distance of each place field to the edge of the chamber in the 1^st^ session, with one group being close to the edge (dashed lines, distance to edge ≤ 3 mm, 233 ± 135 cells, mean ± s.d.) and the other group being close to the center (dotted lines, distance to edge > 3 mm, 352 ± 171 cells, mean ± s.d.).

**Extended Data Figure 10.**
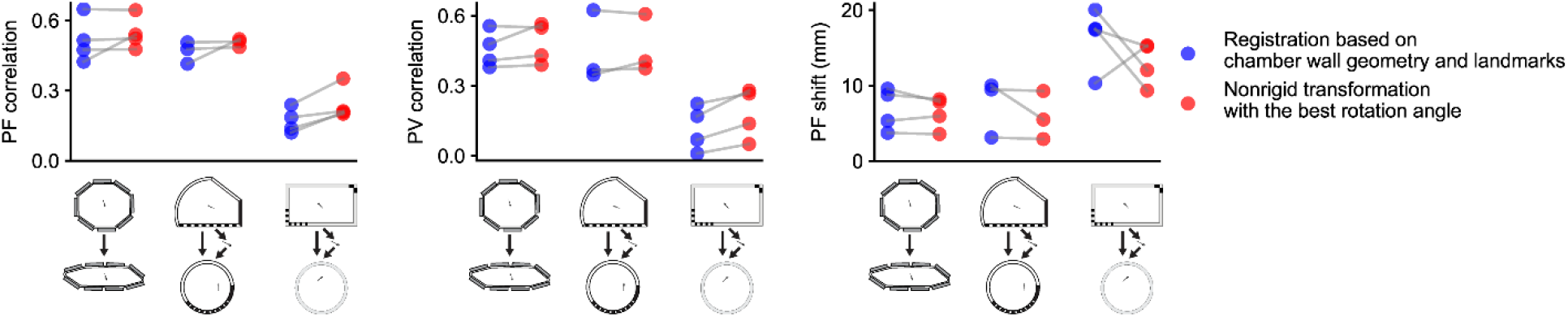
Weakly coherent place field rotations during remapping. The increase in the median PF correlation (left), PV correlation (middle), and PF shift (right) are shown for 3 experiments: wall morphing without fish removal (**Fig. 4o**), wall morphing with fish removal (**Fig. 5i**), and wall morphing with landmark removal and fish removal (**Fig. 5m**). Blue dots show the comparison of session 1 and session 2 after registration of session 2 maps based on session 1 wall geometry and landmarks. Red dots show the comparison of session 1 and session 2 after registering session 2 maps to session 1 maps using the nonrigid transformation with the best rotation angle. Data from the same fish are connected by gray lines. The observed improvement in PF correlation from map rotation was evaluated by a non-parametric shuffle test (**Methods**), applied to each fish individually: wall morphing without fish removal (1/4 fish, p < 10^−5^, 3/4 fish, p ≥ 0.01) wall morphing with fish removal (1/3 fish, p < 10^−5^, 2/3 fish, p ≥ 0.01), and wall morphing with landmark removal and fish removal (2/4 fish, p < 10^−5^, 1/4 fish, p < 0.01, 1/4 fish, p ≥ 0.01).

## SUPPLEMENTARY INFORMATION

**Supplementary Video 1 | Activity of three place cells over the course of one experiment.** Fish trajectory is plotted in gray, and cell activation (Δ*F*(*t*) > 3 s.d.) is shown as colored dots (cyan, magenta, and yellow representing the three cells). Activity is smoothed with a 2.5 s boxcar average filter.

**Supplementary Video 2** | **Activity manifold untangling over time**. We fit an isomap manifold to the last 30 min of imaging session 1 (before chamber rotation) to establish stable axes for 2D embedding and then repeatedly transformed each window of 30 min of population activity data into this embedding (**Methods**).

